# ProteusAI: An Open-Source and User-Friendly Platform for Machine Learning-Guided Protein Design and Engineering

**DOI:** 10.1101/2024.10.01.616114

**Authors:** Jonathan Funk, Laura Machado, Samuel A. Bradley, Marta Napiorkowska, Rodrigo Gallegos-Dextre, Liubov Pashkova, Niklas G. Madsen, Henry Webel, Patrick V. Phaneuf, Timothy P. Jenkins, Carlos G. Acevedo-Rocha

**Affiliations:** The Novo Nordisk Foundation Center for Biosustainability, Technical University of Denmark, Lyngby, Denmark; Department of Biotechnology and Biomedicine, Technical University of Denmark, Lyngby, Denmark

## Abstract

Protein design and engineering are crucial for advancements in biotechnology, medicine, and sustainability. Machine learning (ML) models are used to design or enhance protein properties such as stability, catalytic activity, and selectivity. However, many existing ML tools require specialized expertise or lack open-source availability, limiting broader use and further development. To address this, we developed ProteusAI, a user-friendly and open-source ML platform to streamline protein engineering and design tasks. ProteusAI offers modules to support researchers in various stages of the design-build-test-learn (DBTL) cycle, including protein discovery, structure-based design, zero-shot predictions, and ML-guided directed evolution (MLDE). Our benchmarking results demonstrate ProteusAI’s efficiency in improving proteins and enyzmes within a few DBTL-cycle iterations. ProteusAI democratizes access to ML-guided protein engineering and is freely available for academic and commercial use. Future work aims to expand and integrate novel methods in computational protein and enzyme design to further develop ProteusAI.

## 1 Main

Protein design and engineering (PE) aim to design or optimize proteins for specific purposes, including the design of protein-based therapeutics to improve human health, or the optimization of enzymes for the sustainable production of chemicals, pharmaceuticals, or recycling of plastics and biological waste [1–7]. One of the most successful PE methods is directed evolution (DE), which consists of iterative rounds of gene mutagenesis, screening, and selection [8]. However, DE has a very low success rate and high cost since many rounds of screening are usually needed to sufficiently improve protein function [9]. The problem arises because the vastness and complexity of the sequence space make it difficult to predict which mutations will enhance protein properties [10]. Furthermore, mutations rarely affect a single property; instead, they often result in trade-offs, such as between stability, selectivity, and activity, increasing the complexity even further[11, 12]. Addressing these challenges effectively is crucial for the advancement of PE.

Many key challenges in protein optimization and design are increasingly addressed with ML methods. For example, the discovery of proteins with specific functions can be facilitated by protein language models (pLMs) [13–15]. pLMs capture vast evolutionary information that can be used to predict mutational effects on protein function without requiring experimental data or fine-tuning. Such predictions, refereed to as zero-shot (ZS) predictions, are competitive with supervised mutational effect predictors [16, 17]. Because these models do not require experimental data, they are excellent tools to predict initial experiments. Another useful application of pLMs is the generation of numerical representations of protein sequences, which serve as information-rich inputs for supervised learning algorithms to annotate and discover uncharacterized proteins, or to predict protein properties such as binding, catalytic activity or selectivity. [18– 21].

These surrogate models can be used together with acquisition functions in the BO framework to learn from experimental data and to guide further experiments [19]. The practice of training models from experimental data to predict further experiments is described here as machine learning-guided directed evolution (MLDE), which has been successfully used, for example, to improve human antibodies [22], or plastic-degrading enzymes [23].

Inverse folding (IF) algorithms are a separate class of ML algorithms in enzyme design trained to recover protein sequences given a backbone structure [24–26]. Solving the inverse folding problem yields diverse protein sequences that often exhibit increased thermostability, expression, and solubility [27–29], useful for many applications in biotechnology.

Although many ML methods are open-source, they can be difficult to use for protein engineers who lack the necessary computational skills and expertise. [30]. Examples of powerful yet complex ML tools include ftMLDE [31], DeCOIL [32], and SaprotHub [33]. On the other hand, some user-friendly tools like STAR [34], or ProteinEngine [35] lack open-source availability, restricting their development for future needs.

Conversely, Damietta [36] is openly accessible, but it lacks an open-source framework for community-driven contributions, such as GitHub, thereby limiting future development and customization. Open-source platforms like Rosetta [37], GROMACS [38], and OpenMM [39] have advanced various aspects of computational biology, including protein design and molecular dynamics simulations, exemplifying the benefits of community-driven development and accessibility. Inspired by the success and utility of these long-term projects, we identified a crucial need for ML tools for protein design and engineering that offer user-friendliness and open-source accessibility.

Accordingly, we developed ProteusAI, an open-source ML platform that is accessible as a web app, as a Python package, and as open-source code on GitHub, enabling local installation and community-driven development. ProteusAI offers modules targeting specific challenges, providing a user-friendly and intuitive interface. The modules offer systematic approaches to annotate and discover novel proteins assisted by representation learning algorithms, the possibility to design novel sequences with IF algorithms to increase their stability, expression and solubility, the generation of mutant libraries through ZS predictions, and iterative optimization through BO algorithms. The modules of ProteusAI are inspired by the DBTL-cycle and meant to be intuitive to use for experimental scientists.

ProteusAI is free for academic and commercial use and available as a web-application through *proteusai*.*bio*, as a Python package through PyPI *proteu-sAI*, and as open-source code for local installation on GitHub under *jonfunk21/ProteusAI*. The app and its connection to the DBTL cycle of protein optimization are illustrated in Figure 1.

**Figure 1:**
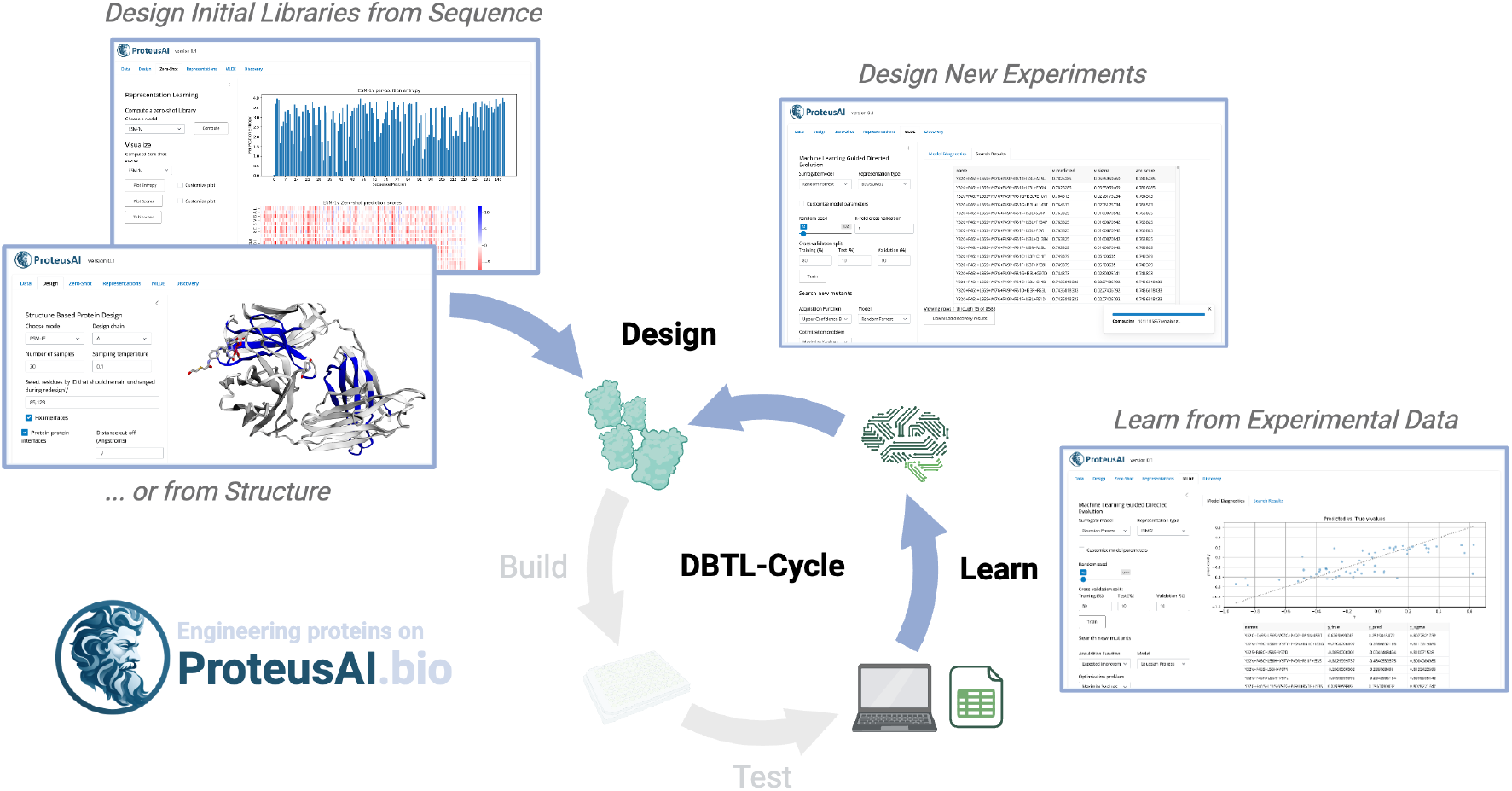
ProteusAI and the DBTL-cycle. The ProteusAI platform is designed to assist protein engineers in the design and learn stages of the Design-Build-Test-Learn (DBTL) cycle. The user-friendly app provides modules to begin a DBTL-cycle given an protein structure or a sequence. The MLDE module of the app is centered around learning from experimental data and the prediction of further experiments.

## 2 Results and Discussion

### 2.1 ProteusAI Modules Overview

ProteusAI provides four modules designed to assist various stages of the DBTL cycle, whether the starting point is an individual protein sequence, structure, or a protein library with or without experimental data. The following section introduces the modules and their intended use in the DBTL cycle. Identifying a functional start sequence is the first step of the DBTL cycle. However, it can be difficult to decide which sequence to start with when presented with a large number of potential sequences than can be tested. The *Discovery module* helps researchers to predict functional clusters of sequences, offering a systematic approach to select diverse yet functionally similar protein sequences.

Once an initial sequence has been identified and tested, other properties such as solubility or stability likely need to be improved for downstream uses or novel applications. The *Protein Design module*, offers structure-based design algorithms that have shown promise in improving these properties. This is also useful, when protein activity and selectivity should be improved via PE or DE [40], which may decrease stability [41].

Finally, MLDE is an effective method to iteratively improve one or more properties, such as catalytic activity or selectivity of enzymes. The *Zero-Shot module* can be used to predict initial mutant libraries to start the collection of experimental data. This data can then be used to train ML models in the *MLDE module* to recommend further experiments. Both the *MLDE-* and the *Discovery-module* rely on numerical representations of proteins that can be computed, and visualized, in the *Representations module*. An overview of the modules is illustrated in Figure 2.

**Figure 2:**
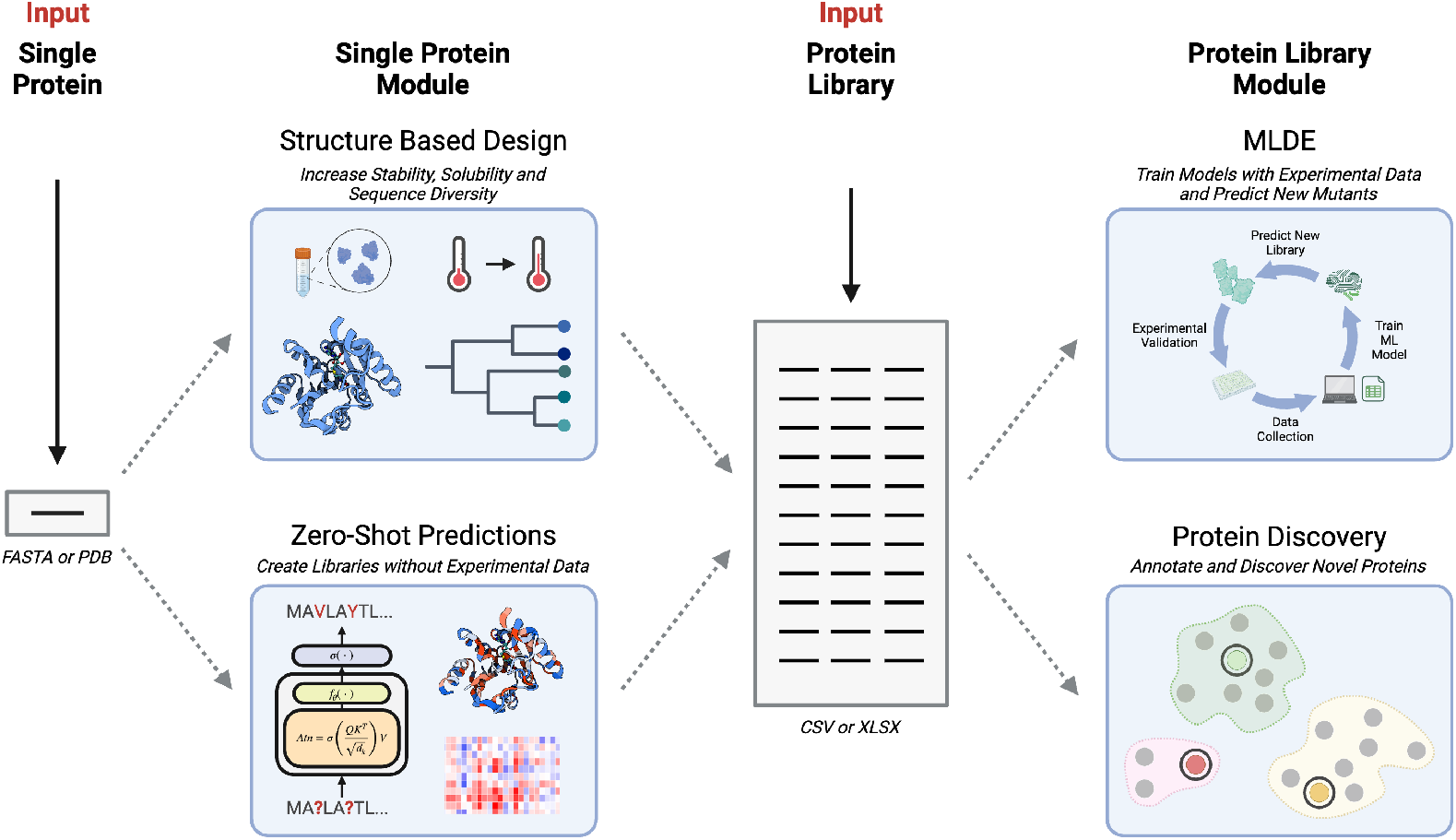
ProteusAI modules. A visual representation of modules that support different stages of workflows in protein engineering and design. Users can either start from a single protein file (sequence or structure) to diversify the protein through structure-based design with IF algorithms or use pLMs to create an initial mutant library through ZS predictions. Users can use the MLDE module to recommend future experiments if experimental data is available. Furthermore, partially annotated data can be used to predict the annotation of unknown proteins, and unannotated data to sample diverse subsets of sequences.

**Figure 3:**
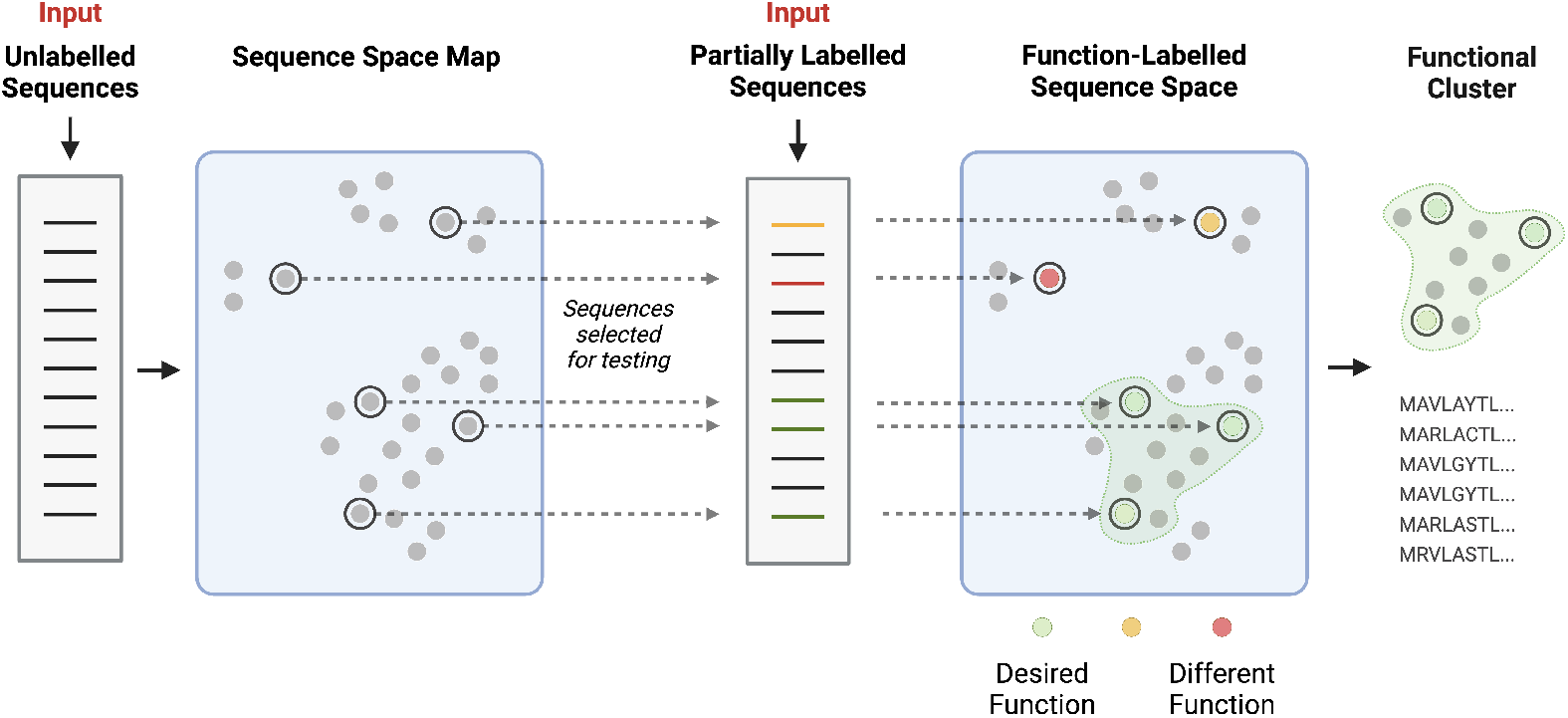
Discovery module. Protein sequences are mapped onto a functional representation space using a pLM. Unlabelled sequences (left panel) are clustered based on inferred functional similarities. Diverse candidates are selected for synthesis and testing, shown as highlighted clusters, where experimental labels are fed back into the module for further refinement. As more functional data is incorporated (right panel), clusters of sequences become annotated with known functional labels, allowing for targeted discovery of novel proteins with desired characteristics. The process helps identify functional candidates with a high probability of functional activity, facilitating a systematic approach to protein discovery.

#### 2.1.1 Protein Discovery Module

PE projects often require searching large sequence databases to discover proteins with specific functions, such as catalytic activity or expression potential in a novel host. However, these searches frequently often yield more candidate sequences than can be practically tested, posing a significant challenge in determining which sequences to prioritize. The ProteusAI Discovery module addresses this issue by leveraging pLMs that map protein sequences onto a high-dimensional representation space, in which sequences are clustered based on functional similarities [13].

Recent studies have shown that pLMs can encode rich structural and functional information, enabling the discovery of proteins with similar or novel functionalities even in the absence of explicit annotations [42]. The *Discovery module* provides a systematic and scalable approach for identifying diverse sequences which are functionally similar.

The module accommodates two different starting points depending on the availability of functional annotations:

##### Unlabelled Data

When starting with sequences lacking functional annotation, such as those identified by homology searches (e.g., BLAST [43]), the *Discovery module* can cluster the candidate sequences and sample diverse sequences from one or many clusters. These selected sequences can then synthesized and tested. Functional data from these experiments can then be used for the next use case.

##### Partially Annotated Data

Once some functional labels are available, the *Discovery module* leverages this information to improve functional prediction, increasing the efficiency of future rounds of sequence selection.

Through this iterative refinement process, the Discovery module seamlessly transitions from handling entirely unlabelled data to leveraging partial annotations, offering a dynamic, data-driven approach to optimize the discovery of functional proteins.

#### 2.1.2 Structure-based Protein Design Module

In PE, it is often necessary to enhance the expression, solubility and stability of proteins before optimizing activity or selectivity. IF algorithms, trained to predict the protein sequences from protein backbone structures, have shown promise to increase the expression, solubility, and thermostability of proteins and enzymes while preserving their native function - if they are constrained from mutating functionally important regions. Examples of these regions are protein-protein, protein-ligand, or metal-ion binding interfaces [44], as well as functionally important catalytic and loop residues, and evolutionary conserved regions [45].

In the *Design module*, IF algorithms can be used and tailored to generate diverse protein sequences from protein backbone structures with the goal of improving properties such as expression, solubility, thermostability, and potentially even catalytic activity, which was observed in diverse protein families [27–29]. Based on these IF studies, we recommend constraining the IF algorithm to preserve functionally important sites and evolutionary conserved residues (Supplementary Table S8). To achieve this goal, the *Design module* offers a tool to automatically detect functionally important sites, such as protein-protein, protein-ligand, and protein-ion interfaces based on atomic distances. Additionally, evolutionarily conserved sites can be entered manually to prevent them from being redesigned. These residues can be determined using co-evolution tools like EVcouplings [46], which are easily accessible online. Sequences sampled with IF algorithms can have low similarity to the original sequences and may serve as novel starting points for MLDE experiments. In addition, preemptively increasing the thermostability using the *Design module* can increase the evolvability of proteins and mitigate trade-off effects [41]. The methods section provides detailed information on the use of IF algorithms in ProteusAI. Figure 4 gives an overview of the *Design module*.

**Figure 4:**
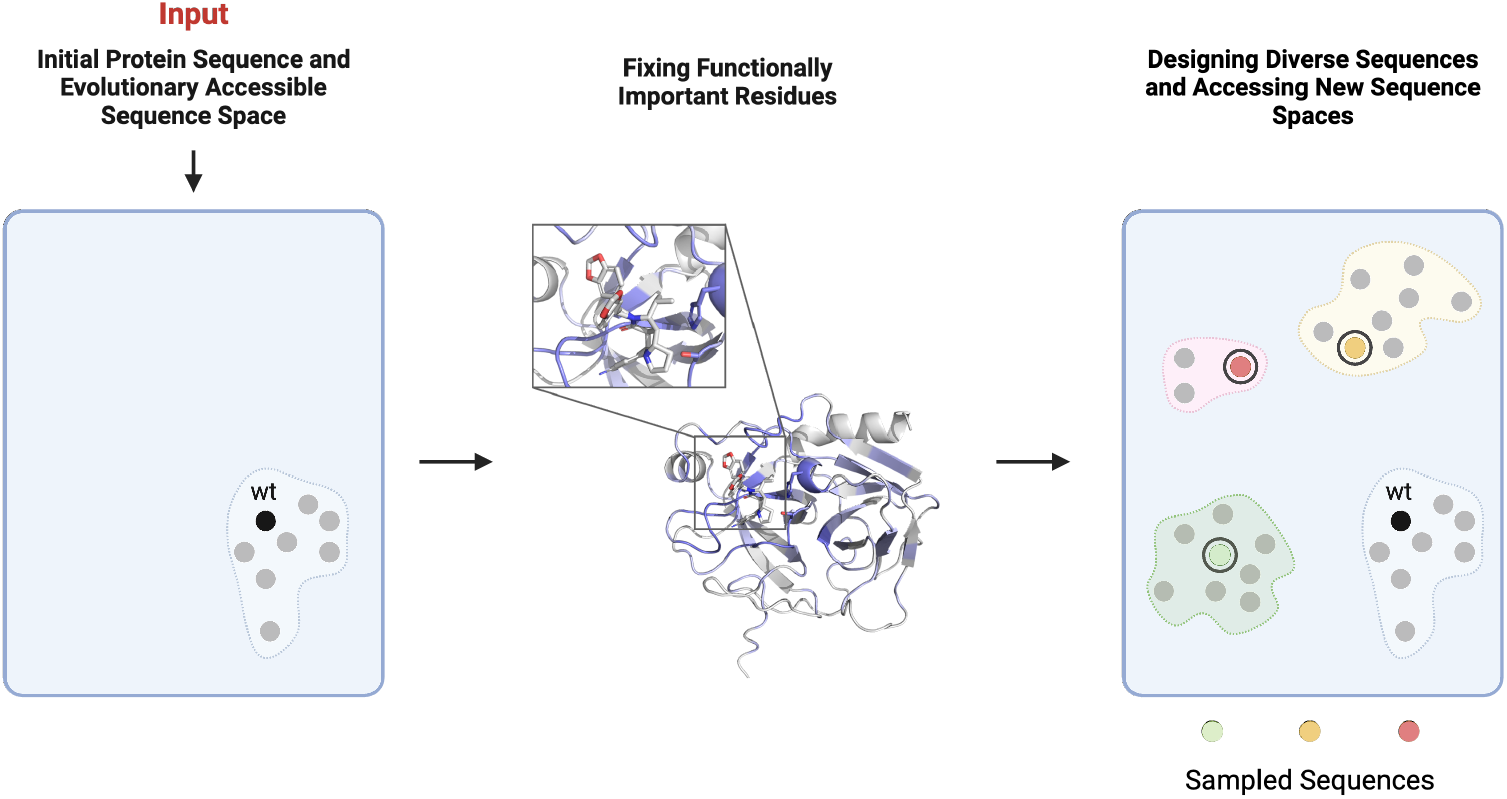
Design module. A visual representation of a wild-type protein and the sequence space that is accessible through directed evolution. In this module, protein sequences are re-designed given an input structure, which can be tailored to preserve functionally important residues. Diverse sequences can then be sampled which are likely to be more stable than the original sequence and distant in sequence space, allowing for further protein optimization.

**Figure 5:**
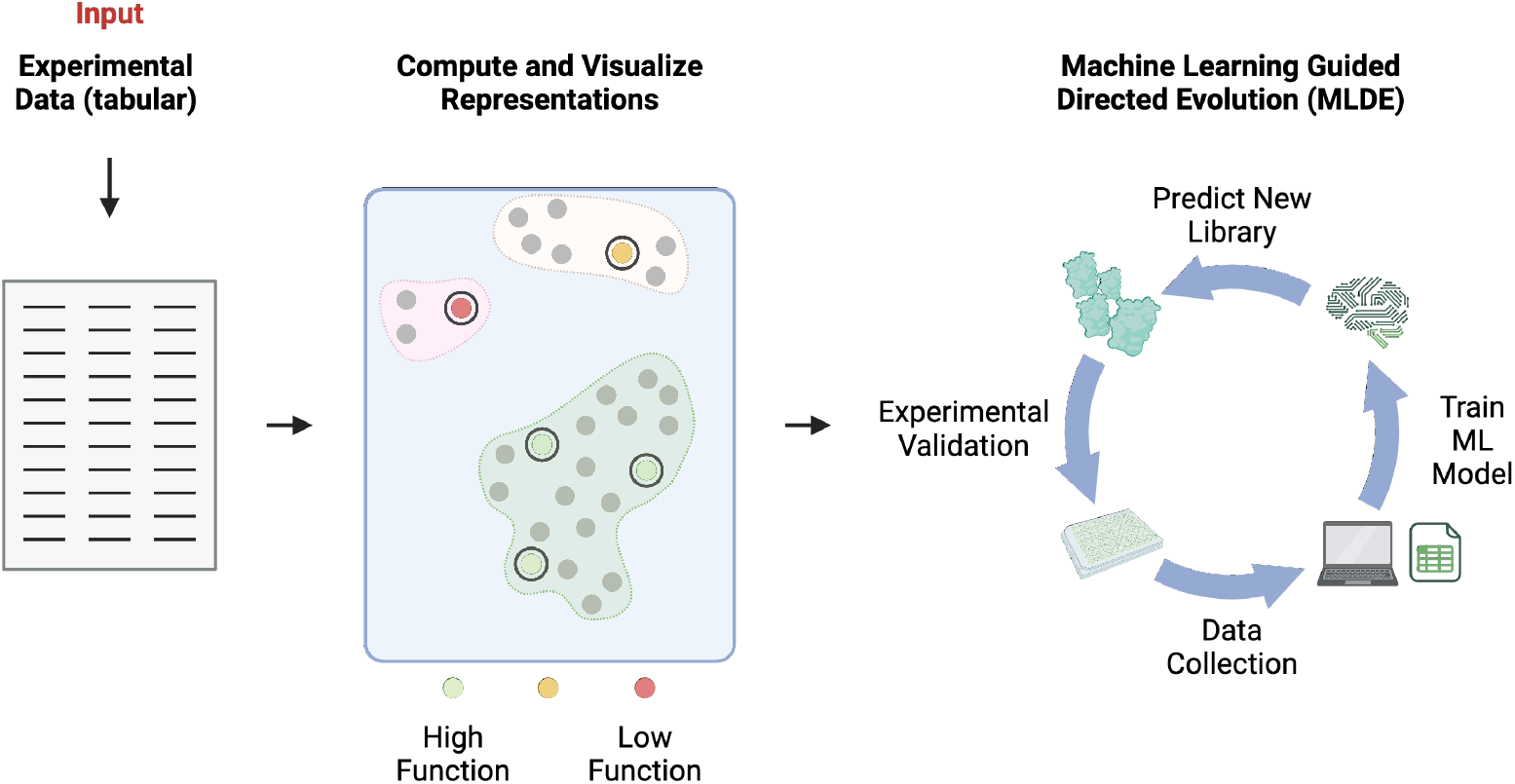
Machine Learning Guided Directed Evolution (MLDE) module. The module starts by uploading experimental data to the app, stored in a tabular format. After uploading the data, the user can compute and visualize protein representations before training ML models to predict future experiments.

#### 2.1.3 Initial Library Design with Zero-Shot Predictions

One of the most effective methods of protein optimization is MLDE, where proteins are optimized through iterative testing, data collection, model training, and prediction of further experiments [19]. MLDE experiments start with an initial mutant library, which can be difficult to generate without prior data. The *Zero-shot module* of ProteusAI is designed to create an initial mutant library using the ZS inference of pLMs to predict mutational effects based on amino acid substitution probabilities [47]. The substitution probabilities result from the masked-language paradigm [48], where models are trained to reconstruct a partially masked and corrupted input sequence. During ZS inference, the model assigns probabilities to all possible amino acids (tokens) for the masked or corrupted positions. The predicted substitution scores correlate with protein fitness, such that high scoring substitutions are often beneficial for fitness and low scoring substitutions detrimental [49]. Because of this correlation, libraries that are generated from high scoring mutants are enriched in functional variants compared to randomly generated libraries (Supplementary Figure S1). In ProteusAI, ZS libraries are created by selecting mutants with the highest the masked marginal probability (MMP) scores explained in the methods section 4.3. These libraries can then be experimentally validated to generate data for further optimization, entering the *MLDE module*. The ZS module uses the pLMs ESM-1v [49] and ESM-2 [13], to generate ZS scores based on Equation 1.

#### 2.1.4 Protein Representation Module

The previous sections described how self-supervised models (pLMs and IF models) can be used to improve proteins, without experimental data. On the other hand, supervised ML models rely on experimental data to predict continuous properties including catalytic activity, stability, binding affinity, or fluorescence (regression), or discrete properties such as the protein family, enzyme commission (EC) number, or protein location (classification) [50]. These supervised ML models use numerical representations of proteins that can be generated with classical algorithms or pre-trained ML models like pLMs. These representations differ in their inductive biases, prior knowledge or assumptions about the data that affecting the training efficiency, and the computation time, which affects the number of sequences that can be evaluated by a model in any given time [20]. These representations can be computed and visualized in the *Representation module* and are used by the *MLDE-* and the *Protein Discovery module*.

The simplest sequence representations that can be generated are one-hot encoding (OHE), where amino acid sequences are converted into matrices, where each column represents an amino acid. Each row represents a position in a sequence, and each row in an OHE matrix contains a single ‘1’ in the column corresponding to the amino acid at that position in the sequence, and all the other values are ‘0’. This form of representing sequences assumes that all amino acids are equally different, thus it has a low inductive bias [20].

BLOSUM representations are alternative, fast-to-compute representations that represent residues by their evolutionary derived BLOSUM substitution scores [51]. The evolutionary inductive bias has been shown to increase the performance of ML models compared to OHE sequences [31].

In contrast, pLMs like ESM-2 [13], ProtGPT2 [15], or ProtTrans [14] excel at extracting representations that are rich in evolutionary information, but are expensive (slow) to compute [20]. pLM representations are highly effective for training surrogate models, often reported to correlate very well with experimental data [47]. Due to their advantages and ease of use, the pLMs used in ProteusAI are ESM-2 [13] and ESM-1v [49].

To summarize, both OHE and BLOSUM encodings are fast to compute, which enables the computation of many mutants, leading to greater exploration of potential search spaces. However, the quick computation comes at the cost of reduced predictive power. The optimal representation for a problem may differ on a case-by-case basis, thus the user has the choice to prioritize search speed by choosing OHE or BLOSUM representations or predictive accuracy by choosing pLM representations.

#### 2.1.5 Machine Learning Guided Directed Evolution (MLDE) Module

MLDE is a powerful strategy to iteratively optimize protein properties such as catalytic activity, binding affinity, and thermostability [19]. By integrating experimental data with advanced ML methods, the *MLDE module* provides a systematic approach to enhance protein function.

The core of the *MLDE module* are powerful Bayesian Optimization (BO) algorithms, which are increasing used for applications in PE [52]. BO algorithms consist of two key components: a probabilistic surrogate model (SM) that predicts fitness values and uncertainties, and an acquisition function (AF) that weighs the prediction and uncertainty to prioritize novel sequences for further testing [53, 54]. This combination enables balancing exploration and exploitation and the operation in confidence regions of the fitness landscape.

The choice of representation and model type are critical, as it impacts both the prediction accuracy and the computational efficiency of the process. Accurate models allow for better differentiation between improved and non-improved mutants. At the same time, efficient computation enables the evaluation of more sequences *in silico* before proceeding with experimental validation.

The AF plays a pivotal role in guiding the selection of sequences for future experiments. By balancing exploration (evaluating uncertain areas of the sequence space) and exploitation (focusing on sequences predicted to yield high fitness), the AF ensures a comprehensive search of the sequence space. The Upper Confidence Bound (UCB) function is the most optimistic acquisition function, exploring sequences with high potential despite higher uncertainty, whereas the Expected Improvement (EI) will penalize high uncertainties, striking a balance between exploration and exploitation. The greedy acquisition ignores uncertainty values.

The MLDE module uses a genetic algorithm (GA) to propose mirroring experimental DE methods [9]. The GA prioritizes the combination of beneficial mutations before considering random mutations; however, the preference can be tailored to include more or fewer random mutations. The algorithm is explained in detail in the Supplementary Information (Alg. 2). The performance of the MLDE module is benchmarked in Section 2.2. Based on resulting data illustrated in Section 2.2, we recommend using a Random Forest (RF) model with either ESM-2 or BLOSUM62 representations. RFs trained on ESM-2 representations are more accurate but significantly slower.

### 2.2 MLDE Benchmarking

Optimizing proteins with the lowest number of experimental rounds along the Design-Build-Test-Learn (DBTL) cycle is a critical goal for wet lab scientists. With limited initial data and the high costs associated with each round of experimentation, selecting the most efficient models can make a significant impact on the success of PE projects. The benchmarks presented here aim to provide insight into which ML models and acquisition strategies are most effective at guiding experimentalists in selecting optimal mutants with minimal experimental effort.

We focus not only on traditional evaluation metrics like mean-squared error (MSE), but also on metrics that reflect real-world outcomes, such as the number of DBTL rounds needed to identify the best possible mutants. By simulating DBTL cycles on various published datasets, we provide a comprehensive assessment of different model performances across multiple sample sizes and protein properties, which are key considerations especially for users navigating protein landscapes with low-data settings.

#### 2.2.1 Model Efficiency and Sample Size Impact

As shown in Fig. 6a, the RF model trained on ESM-2 representations and coupled with the EI AF consistently outperformed other models, needing fewer DBTL rounds to identify the top N mutants. However, smaller sample sizes (5 sequences per iteration) sometimes led to model failures to discover the top mutants, as indicated by the grey stars in Fig. 6a. This underscores the importance of data availability in model-driven PE tasks, where even small increments in sample size can lead to significant reductions in experimental workload.

**Figure 6:**
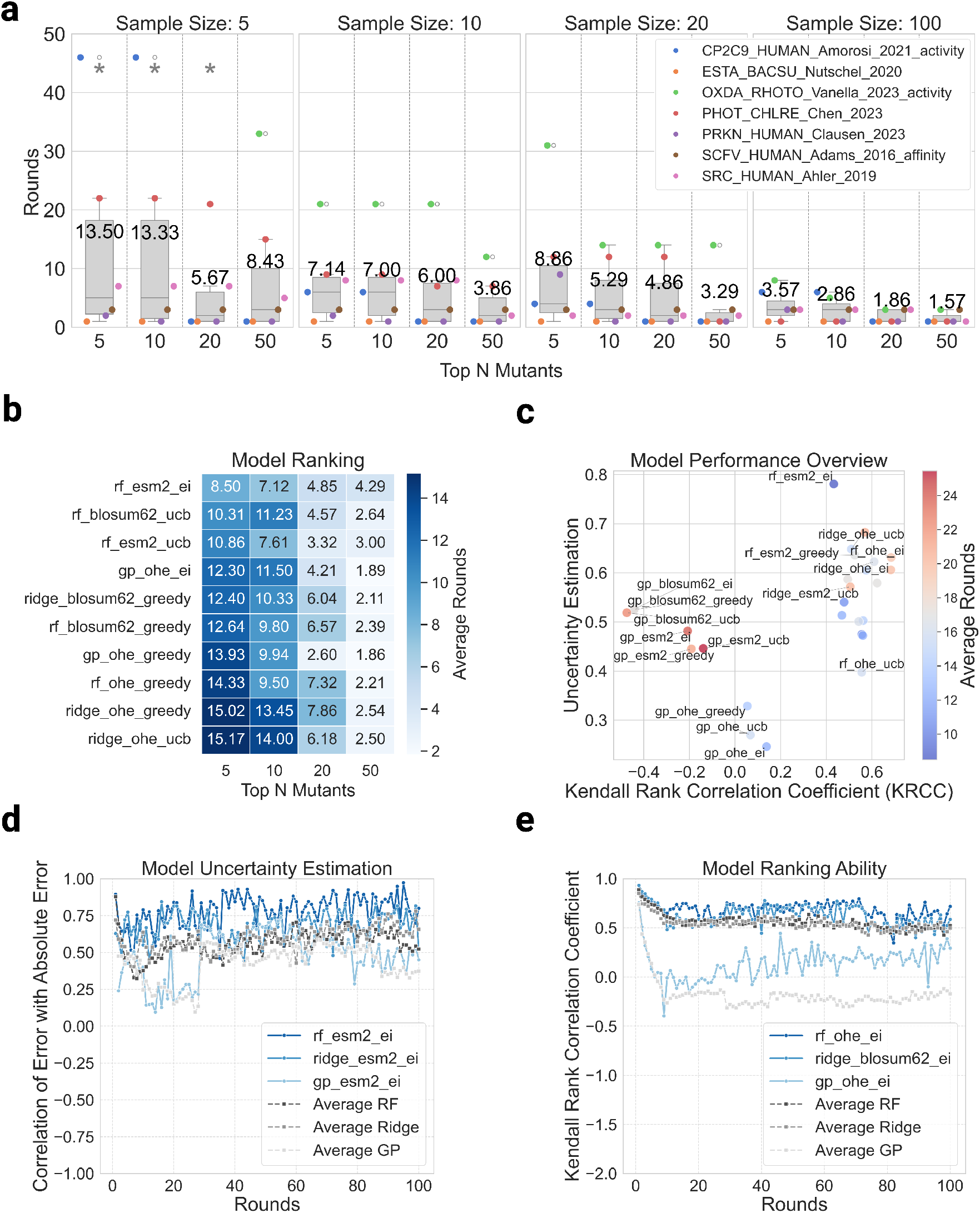
Benchmarking overview. Summary of the efficiency of various models and acquisition strategies in discovering top-performing protein mutants across multiple datasets and sample sizes. **(a)** Box plots show the number of DBTL rounds required to identify the top N variants (5, 10, 20, 50) across sample sizes of 5, 10, 20, and 100. The RF model trained on ESM-2 representations with the EI acquisition function consistently discovered top mutants with the fewest rounds, especially for larger sample sizes. Grey stars indicate datasets where top variants were not discovered within the 100-cycle limit. **(b)** Heatmap ranking models by the average number of rounds required to discover top mutants across all datasets and sample sizes. RF models, particularly with ESM-2 and BLOSUM62 representations, outperform GP and Ridge Regression (RR) models, especially for smaller variant sets (Top 5 and Top 10). **(c)** Scatter plot of KRCC against uncertainty error (UE), colored by the number of rounds required to discover top mutants. High KRCC and UE values are generally associated with better model performance, though outliers are present. **(d)** Line plot showing the correlation between model uncertainty and absolute error (uncertainty estimate - UE) as a function of DBTL rounds. While RF and RR models maintain relatively stable uncertainty estimates, GP models show significant fluctuations, improving only after more data is collected. **(e)** Line plot of the KRCC over DBTL rounds, indicating that RF and RR models maintain a high ability to rank proteins correctly throughout the process, while GP models improve after gathering sufficient data.

For practitioners, this insight suggests that while RF models are well-suited to low-data regimes, the risk of failure with smaller sample sizes needs to be carefully considered. Increasing the initial data volume may help avoid unnecessary rounds if possible.

#### 2.2.2 Model Comparison Across Datasets and Acquisition Strategies

Figure 6b shows clear distinctions in the performance of RF models compared to Gaussian Process (GP) and RR models. RF models, particularly those using ESM-2 representations and the EI AF, were most effective at identifying top variants across different sample sizes. GP models, although traditionally favored for their uncertainty quantification in BO, struggled with smaller datasets but improved their performance over time with more data. This finding has direct implications for users: in data-scarce environments, tree-based methods like RF may provide more reliable results than GP models. However, as datasets grow larger, GPs can become competitive and may offer advantages in uncertainty estimation that are beneficial in later stages of the DBTL cycle.

#### 2.2.3 Uncertainty Estimation and Predictive Power

Figure 6c-e provide further insights into model performance by analyzing how models’ uncertainty estimation and ranking ability evolve with DBTL rounds. In particular, Fig. 6c shows that models with higher Kendall rank correlation coefficient (KRCC) [55] and robust uncertainty estimates (UE) tend to perform better at identifying top mutants, hinting at the importance of ranking-based metrics in guiding experimental design. The correlation between high KRCC and UE, and model success, suggests that models capable of correctly ranking protein variants are effective in driving the discovery of optimal candidates. For practical users, focusing on these diagnostics during model selection could significantly reduce the number of experimental rounds required to find the best candidates, streamlining the PE process.

#### 2.2.4 Performance Stability Across DBTL Rounds

In Fig. 6d-e, RF models exhibit relatively stable uncertainty estimates and ranking ability (high KRCC) throughout the DBTL process. GP models, by contrast, show greater fluctuation in early rounds but improve as more data becomes available. This pattern suggests that GP models, while suitable for datasets with abundant data, may not be as effective in early-stage discovery tasks where data is limited. Conversely, RF models offer a more consistent performance, making them a better choice when starting with smaller datasets.

#### 2.2.5 Practical Implications for Protein Engineers

The benchmarks presented here provide key insights into the trade-offs between different models and acquisition strategies in the DBTL cycle. In scenarios with limited data, which is often the case during early-stage experimental design, RF models combined with the EI AF show consistent success in reducing DBTL iteration steps. For protein engineers aiming to streamline their workflows, adopting models that perform well in low-data environments can minimize costly experimental iterations and accelerate the discovery of top-performing mutants.

However, the choice of model should also take into account dataset size and characteristics. While GP models become more competitive with larger datasets, they may not be the optimal choice for initial stages of the DBTL cycle. Future work could explore hybrid strategies that combine the strengths of tree-based models and GP uncertainty estimation for broader applicability in PE and MLDE workflows.

## 3 Conclusion

ProteusAI represents a novel resource for ML-guided protein engineering and design by providing an opensource and user-friendly platform that integrates state-of-the-art ML methods into the DBTL cycle. Through its modular approach, researchers can seamlessly explore protein discovery, structure-based design, zero-shot predictions, and iterative optimization using MLDE. Our benchmarking results demonstrate that ProteusAI effectively accelerates the identification of top-performing mutants, particularly when leveraging RF models with ESM-2 representations and the EI acquisition function.

The study also reveals important insights into model performance in low-data regimes, emphasizing the value of metrics such as the KRCC and UE in predicting success. While GP models offer robust uncertainty estimates, they require larger datasets to perform optimally, positioning RF models as more effective in smaller sample-size settings. Finally, these findings underscore the need to consider trade-offs between experimental volume and the number of rounds needed for optimization.

In future work, we aim to reduce the computational cost for generating pLM representations and explore novel methods that could further enhance ProteusAI’s impact. As we continue to refine our platform, ProteusAI will serve as a key tool in democratizing and accelerating ML-guided protein engineering.

## 4 Methods

### 4.1 Discovery Module Methods

The discovery module aims to sample diverse sequences yet functionally similar sequences. The function prediction can be generated either from unsupervised clustering algorithms or supervised classification algorithms. Once sequences have been labeled, the GA (Alg. 1) can be used to sample diverse sequences from selected clusters. The GA used in this module aims to increase the euclidean distance between the sampled representations, leading to more diverse samples with the same predicted function.

#### Algorithm 1

Sampling Diverse Sequences.

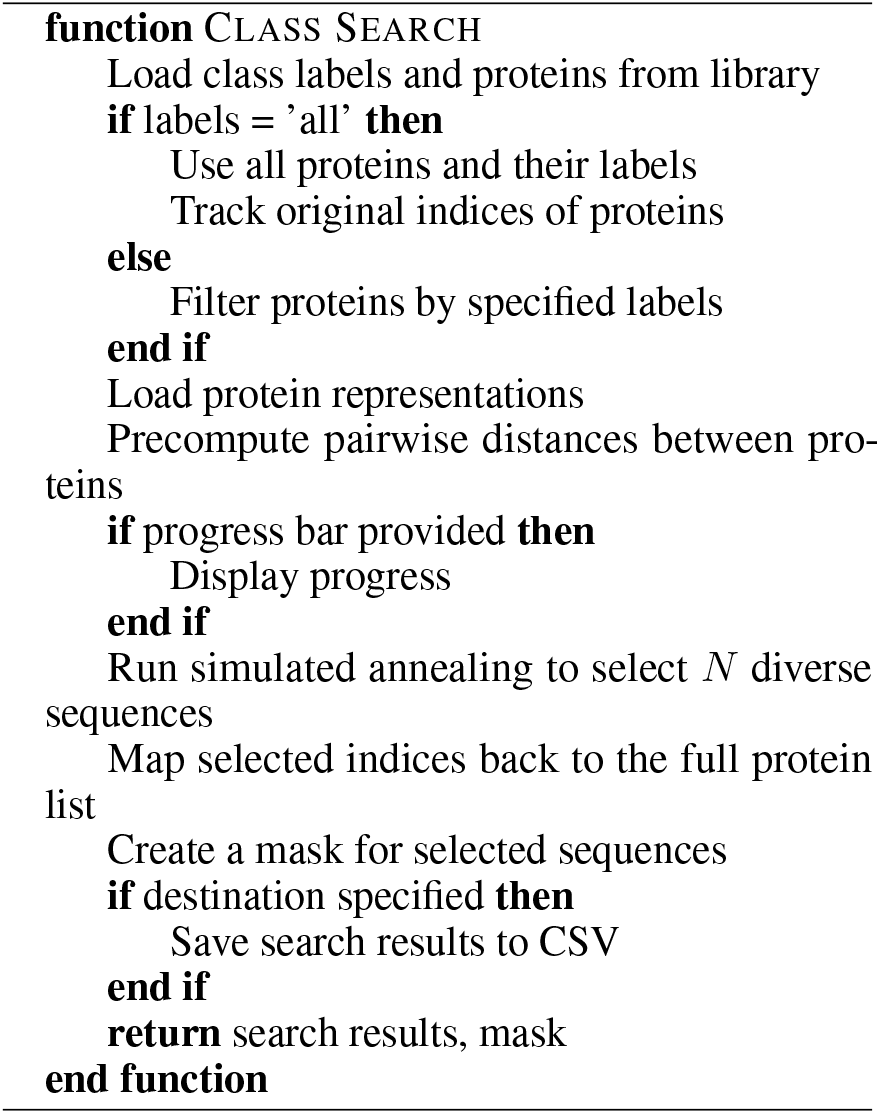

### 4.2 Design Module Methods

The Design Module uses the inverse folding algorithm ESM-IF and ProteinMPNN to sample sequences given an input backbone structure [25, 26]. The geometric vector perceptron (GVP) used for ESM-IF incorporates invariant geometric processing layers to handle structural input. It uses an autoregressive framework to perform inverse folding, predicting sequences based on the spatial coordinates of backbone atoms (N, C*α*, C) without explicit consideration of side chains. On the other hand, ProteinMPNN operates by leveraging a message-passing neural network (MPNN) that processes structural features, including interatomic distances and backbone dihedral angles. The model also uses an autoregressive decoding approach to predict amino acid sequences at each position in the backbone structure. In ProteusAI, the user has the option to control the number of samples that should be generated by the models, the protein chain that should be sampled for (for multi-chain inputs), and the sampling temperature. Based on previous studies listed in the supporting Table S8 we recommend to sample thousands of sequences, using different temperatures (e.g. 0.01, 0.1, 0.3) followed by rigorous filtering. For filtering we recommend using folding models, such as ColabFold [56], which is an openly accessible, easy to use implementation of AlphaFold 2 [57], to filter out low confidence predictions and structures that include heavy atom clashes. Furthermore, we recommend a thorough literature research to identify functionally important residues in the structure to fix them during the re-design process. The sampling temperature can be tuned to sample more or less diverse sequences.

### 4.3 Zero-Shot Module Methods

The *ZS module* uses pLMs to predict mutational effects on protein fitness without prior data. These predictions can be used to create an initial mutant library for MLDE experiments. ZS scores are computed using the masked marginal probability (MMP), proposed by Meier et al. [49] described in equation 1.

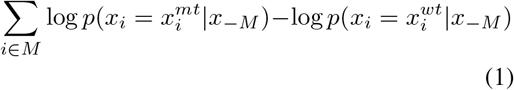

The recommended library is constructed by sorting the MMP values from the highest to the lowest values and selecting the highest-scored mutant for N positions, specified by the user. Alternatively, the ZS predictions can be downloaded as a CSV file. It is recommended to use ESM-1v to compute ZS scores, which have been shown to correlate better with experimental results than ESM-2 [58].

### 4.4 Representations Module Methods

In the *Representations module* pLM representations can be computed from the models ESM-2 and ESM-1v, or classical algorithms yielding OHE or BLO-SUM representations. The pLM representations are computed by forward passing sequences through the pLM extracting the final-layer output tensor. The classification token is then removed from the tensor, and the tensor averaged over the sequence length dimension, yielding the representation. For supervised learning, we recommended to use ESM-2 over ESM-1v, because, because predictions of models trained on ESM-2 correlate better with experimental data [58].

### 4.5 MLDE Module Methods

The *MLDE module* builds on experimental data, to predict further experiments. The library data used for this module are tables of protein sequences with associated fitness values, and descriptive names. ProteusAI expects the input data to be in the format of an Excel file or a CSV file, that include a column for the protein sequences, a description of the mutant (e.g. A17V), and a column for protein fitness. For more information on the data format and recommended normalization steps, please refer to the supporting information sections 9.1 and 9.1.

The core of the MLDE module is the BO loop, which consists of probabilistic surrogate models that are trained to map protein representations to fitness values, a genetic algorithm to propose novel sequences, and an acquisition function rank the sequences for experimental validation. The current available elements for the BO loop are listed in the supporting information Table S2.

#### 4.5.1 MLDE Model Diagnostics

Diagnosing model performance after training is an essential step for practitioners of machine learning. A key diagnostic is the scatter plot of predicted y-values versus true y-values, where,ideally all points should align along the diagonal, indicating perfect predictions. In reality, this plot helps to visually assess deviations and can reveal areas where the model performs poorly. Accompanying this visual, the *R*^2^ value provides a quantitative measure of how well the model explains variance in the data, with values closer to 1 indicating better performance. A common to look out for, especially when training only on few data points, is that the model may simply predict the same value for each mutant, indicated by a flat line in the diagnostics plot, or same prediction values in the validation table. Here we recommend either trying the stratified split method to split data into train, test and validation datasets, that aims to equally sample sequences from different y value ranges. If the problem persists, one should consider to balance the dataset manually, for example by removing some sequences that have over represented y values, or collecting more data before proceeding.

In addition to visual diagnostics, summary statistics play a critical role in evaluating model behavior. In this study, the KRCC is used to assess the ordinal relationship between predicted and true rankings, providing a robust measure of association even when the relationship is not strictly linear. Furthermore, the correlation between model uncertainty and absolute error is another crucial metric, as it reveals how well the model’s uncertainty estimates reflect actual errors. A strong correlation suggests that the model appropriately quantifies its confidence in predictions.

Finally, standard error metrics such as Mean Absolute Error (MAE), Root Mean Squared Error (RMSE) or Pearson Correlation Coefficient offer concise numerical insights into overall prediction accuracy, allowing for comparative evaluations across datasets or models.

#### 4.5.2 MLDE Genetic Algorithm

In the BO loop, a GA is employed to propose novel protein sequences for evaluation. This algorithm is inspired by experimental MLDE approaches, where new mutations are preferably generated by combining mutations that have lead to improvements, together with some random mutations. The resulting mutant sequences are then evaluated by the model, and the process is repeated to optimize the search for improved proteins. In ProteusAI the user can adjust the exploration of random mutations with the exploitation of already known mutations with the ‘explore’ parameter, which ranges from 0 (very exploitative) to 1 (very exploratory). The algorithm won’t create duplications of sequences, thus even very exploitative settings will approach explorative algorithm if the combinatorial space of known sequences should become exhausted.

##### Algorithm 2

Discovering Novel Mutations Using MLDE.

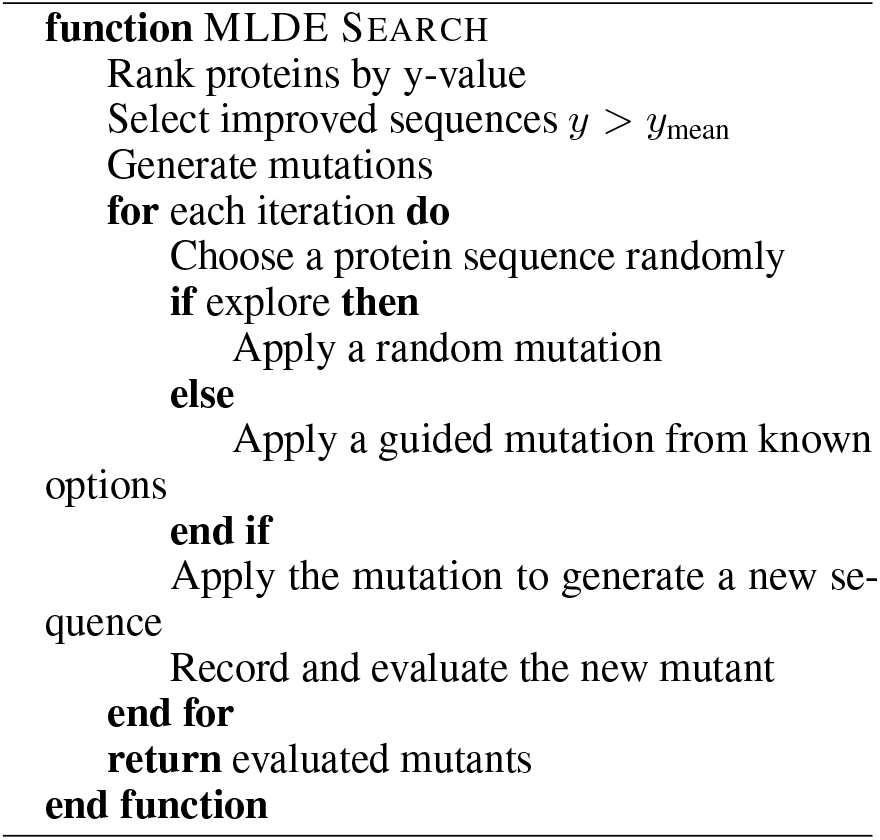

#### 4.5.3 MLDE Benchmarking

In order to assess model performance in the context of protein engineering, we designed a comprehensive benchmarking framework focused on evaluating ML models across multiple datasets and simulation scenarios. The primary goal of our benchmarks was to determine which models and acquisition strategies best minimize the number of iterations required to discover top-performing protein variants in the Design-Build-Test-Learn (DBTL) cycle.

We simulated DBTL rounds by selecting and evaluating protein mutants from a set of published datasets, starting with a small library of sequences. The highest-ranking protein variants were chosen based on initial ZS predictions and used to train BO algorithms. At each iteration, the models proposed the next sequences for evaluation based on their acquisition strategies. This process was repeated until one of the top five highest-ranking mutants was discovered or a maximum of 100 iterations had been completed.

We only selected mutants that were present in the datasets, meaning no de novo designs were proposed. As a result, the benchmarks focus on ranking and selecting from existing sequences in the dataset rather than creating new ones. This allows us to assess how well models can guide experimentalists in selecting optimal candidates with minimal experimental effort.

#### 4.5.4 Datasets and Properties

We selected a diverse set of benchmark datasets that encompass a range of properties relevant to protein engineering. These include:

- **Binding Affinity:** For evaluating how well models can predict interactions between proteins and ligands or other molecules.
- **Catalytic Activity:** To assess a model’s ability to rank variants based on their enzymatic activity.
- **Thermostability:** A critical property in protein engineering, particularly for industrial applications where proteins must maintain function at elevated temperatures.
- **Fluorescence:** Useful for applications in imaging and sensor development.

Most datasets were sourced from the Prote-inGym benchmark suite [58], which provides accurate, experimentally validated measurements of protein properties. An exception is the SCFV_HUMAN_Adams_2016_affinity dataset, which was sourced from Adams et al.[59], specifically chosen for its high-quality binding affinity measurements.

A summary of the datasets and their properties is provided in supporting Table S7. This variety of datasets ensures that our benchmarks capture a broad spectrum of protein engineering tasks, providing practitioners with insights into model performance across multiple domains.

#### 4.5.5 Model and Acquisition Function Selection

We benchmarked three key models widely used in protein engineering and machine learning-driven experimental design:

- **RF:** Known for its ability to handle small datasets and capture nonlinear relationships, making it ideal for early-stage protein engineering tasks.
- **Gaussian Process (GP):** A popular model for Bayesian optimization due to its uncertainty estimation capabilities. GPs are well-suited for tasks where quantifying prediction uncertainty is crucial for decisionmaking.
- **Ridge Regression (RR):** A baseline linear model that allows us to compare nonlinear models like RF and GP against simpler approaches.

Each model was evaluated using the EI acquisition function, a commonly used strategy in Bayesian optimization that balances exploration and exploitation by selecting sequences with the highest predicted improvement over the current best result.

#### 4.5.6 DBTL Simulation Protocol

Our simulation of the DBTL cycle mirrors the workflow followed by experimentalists who is using ProteusAI:

- **Initial Sequence Library:** The DBTL cycle begins with a small library generated by sampling mutants with the MMP ZS scores generated from the model ESM-1v. For sample sizes < 15 the initial number of sequences was set to 15 sequences.
- **Bayesian Optimization Loop:** After ZS predictions, the selected top mutants are used to train a machine learning model (RF, GP, or RR). Based on the model’s predictions, the next round of sequences is proposed using the acquisition function.

Round Completion: The DBTL round is repeated until one of the top five mutants is discovered or 100 iterations are reached. The rounds completion is measured by counting the number of iterations required to identify top mutants, which serves as a key performance metric

#### 4.5.7 Sampling Sizes and Experimental Throughput

To reflect varying experimental throughput capacities in real-world protein engineering tasks, we simulated different sample sizes for each DBTL iteration: 5, 10, 20, and 100 sequences. This allows us to evaluate model performance under conditions of varying data availability and provides insights into how models scale with increasing data.

Smaller sample sizes (e.g., 5 mutants per iteration) are representative of early-stage, low-throughput experiments where data is limited, while larger sample sizes (e.g., 100 mutants per iteration) simulate higher-throughput settings, which are more feasible in later stages of protein development.

#### 4.5.8 Evaluation Metrics

We used a combination of traditional and ranking-based metrics to evaluate model performance:

- **Mean-Squared Error (MSE):** A commonly used metric that evaluates the accuracy of the model’s predictions against true values. While useful, MSE may not fully capture the practical needs of protein engineering, where ranking and discovering the top variants is often more critical than minimizing prediction error.
- **Kendall Rank Correlation Coefficient (KRCC):** KRCC was particularly important in our benchmarks as it measures the model’s ability to rank variants correctly. This metric reflects the practical goal of guiding experimentalists toward the best candidates for further validation. In protein engineering, relative ranking is often more important than the exact prediction of quantitative values, making KRCC a critical measure for model success.
- **Uncertainty Estimation (UE):** We also evaluated models based on their ability to estimate uncertainty in their predictions. Accurate uncertainty estimation is important for guiding experimental design in cases where data is scarce, and incorrect predictions can lead to costly experimental rounds.

#### 4.5.9 Statistical Analysis

We performed statistical analyses to compare model performance across different datasets and sampling sizes. Box plots were used to visualize the distribution of DBTL rounds required to identify top-performing variants. Heatmaps were also generated to rank models based on their average number of rounds across all datasets and sampling sizes. Additionally, scatter plots were used to correlate KRCC with model UE, providing further insights into the relationship between ranking accuracy and predictive performance.

### 4.6 ProteusAI Python Package and App

The Package of ProteusAI is aimed at super-users who want more flexibility than provided in the app and integration with other methods not currently supported, and at developers who want to test new features. The ProteusAI package provides a powerful and modular framework that makes it easy to handle common data types used in protein engineering, combined with a modular software architecture to interface with ML models. The architecture of the software relies on three primary objects, the Protein-, Library-, and Model-objects, which are specialized on different data types and interface with different models. The Protein object is specialized to handle information associated to single proteins, such as the protein sequence, structure or fitness values. This object offers interfaces to models that operate on single proteins, such as ZS models or IF algorithms. The Library object orchestrates collections of proteins and is the cornerstone for visualizations of libraries and the primary input for the Model object. The Model object takes an Library object as input to fit machine learning models to the protein libraries, offering easy interfaces to surrogate models, acquisition functions, and BO loops. All code is open source and continuous integration ensures basic testing. Documentation for the library can be found on proteusai.readthedocs.io. The ProteusAI app is implemented in Shiny for Python [60]. The underlying ML algorithms are implemented using scikit-learn and PyTorch, and the functionality is packaged into an installable library from PyPI. The app and package are available on GitHub: jonfunk/ProteusAI.

### 4.7 Web Server

The application was deployed with Shiny Server v1.5.22.1017 on a Microsoft Azure Virtual Machine with Ubuntu 20.04 as the operating system.

## 5 Acknowledgements

We thank Miguel González Duque and the research group of Wouter Boomsma for insightful discussions about Bayesian optimization. J.F. thanks Jinbei Li, Lei Yang, and Emre Özdemir for discussions about the MLDE module. J.F. thanks the Novo Nordisk Foundation Center for Biosustainability (NNF20CC0035580) who funded this work. CGAR acknowledges support from the Novo Nordisk Foundation grant number NNF20CC0035580. NGM acknowledges support from the Danish Data Science Academy, which is funded by the Novo Nordisk Foundation (NNF21SA0069429) and VIL-LUM FONDEN (40516).

## 6 Author Information

### Contributions

J.F. designed and implemented ProteusAI, ran all the experiments and analyzed the results. L.M. helped with the BO in the MLDE module, S.A.B. helped with the design of the discovery module, M.N. helped with the selection of the datasets for benchmarking. L.P., R.G.D. and L.M. helped with the web server implementation, P.V.P. advised on user experience improvements of the app. N.G. helped with the ZeroShot and Design module. H.W. helped with the publication of the python package and the package documentation. C.G.A.R. and T.P.J. supervised the project. All authors contributed to the writing of the paper.

## Corresponding Authors

Jonathan Funk, Timothy P. Jenkins and Carlos G. Acevedo-Rocha

## 7 Code and Data Availability

ProteusAI is available through http://proteusai.bio/ and as open-source code for local installation on GitHub https://github.com/jonfunk21/ProteusAI. ProteusAI is free for academic and commercial use.

## 9 Supplementary Information

### 9.1 Data Formats

Each module of ProteusAI expects data in specific file formats. The expected file formats for each module are described in the following Table S1:

**Table S1:** Expected data formats for the individual ProteusAI modules.

| Module | FASTA | PDB | CSV | XLSX |
| --- | --- | --- | --- | --- |
| Discovery |  |  | ✓ | ✓ |
| Design |  | ✓ |  |  |
| Zero-Shot | ✓ | ✓ |  |  |
| Representations | ✓ | ✓ | ✓ | ✓ |
| MLDE |  |  | ✓ | ✓ |

#### Data Convention

In addition to expected data formats, ProteusAI follows conventions for describing experimental data. We recommend adherence to these conventions to ensure compatibility with conditions used in the benchmark studies. Experimental data in ProteusAI should be stored as follows in tabular formats, either as a CSV or XLSX file. The tables should contain the following columns:

1. **Sequence** column: containing the protein sequences.
2. **Description** column: containing mutant descriptions.
3. **Data** column: containing experimental values (Y-values).

The protein sequences column should contain the full (mutated) protein sequence as an amino acid sequence. The mutant descriptions should follow the following format: [wt amino acid][position][mutated amino acid], for example, the mutant A15I would describe a mutation from an alanine to an isoleucine at position 15 in the sequence. Higher order mutants should be entered as the combination of single mutant descriptions separated by a ‘+’ or ‘:’ symbol, e.g. A15I+V132L or A15I:V132L, would describe a double mutant from alanine to isoleucine at position 15 and a mutation from valine to leucine in position 132. The experimental values should follow the following convention: The fitness of the wild-type should be defined as 0, and other sequences should be reported relative to the wild-type, where positive values are improvements in fitness, while negative values are decreases in fitness. For categorical Y-values, such as protein classes for the discovery or annotation of proteins, provide the class labels either as strings or numbers and ensure specify the data type as ‘categorical’ in the app.

### 9.2 Zero-Shot Based Libraries Versus Randomly Sampled Libraries

**Figure S1:**
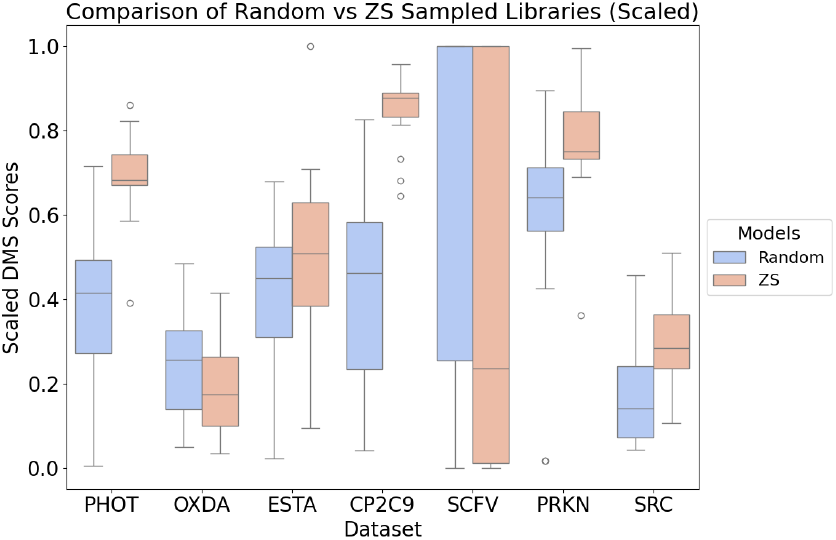
Zero-shot-based libraries have significantly improved fitness over randomly sampled libraries.

### 9.3 Effects of Sample Size

Figure S2 shows the effect of increasing the sample size on the average number of rounds needed to discover the top performing proteins.

**Figure S2:**
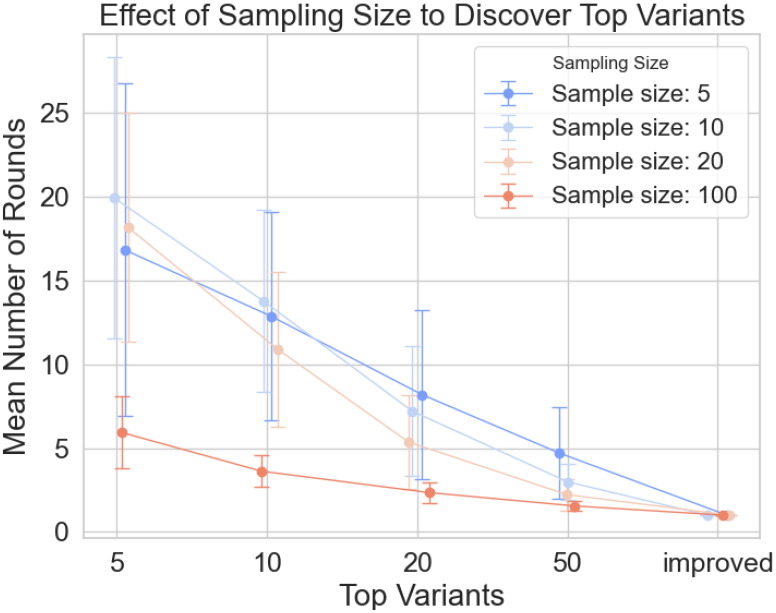
Effect of sample size on number of experiments needed to discover top mutants.

### 9.4 Available Models, Representations and Acquisition Function in ProteusAI version 1.0

The following models, representations and acquisition functions liste in Table S2 are available in ProteusAI version 1.0. They can be expanded in the future to cover more methods.

**Table S2:**
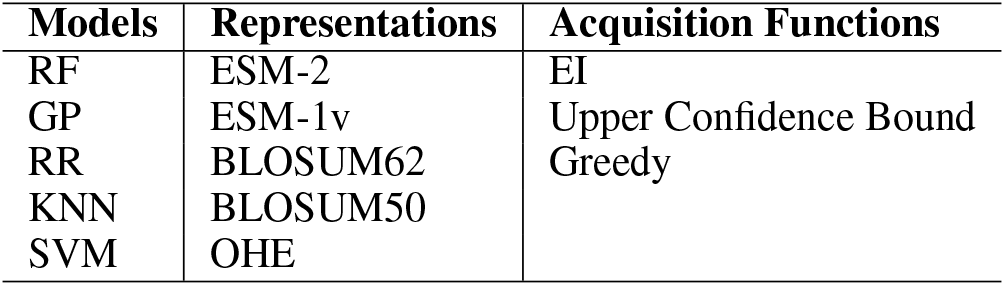
Models, Representations, and Acquisition Functions used in the study.

| Models | Representations | Acquisition Functions |
| --- | --- | --- |
| RF | ESM-2 | EI |
| GP | ESM-1v | Upper Confidence Bound |
| RR | BLOSUM62 | Greedy |
| KNN | BLOSUM50 |  |
| SVM | OHE |  |

### 9.5 Average Number of Rounds to Discover Top N Mutants

The number average number of rounds needed to discover the top variants of every dataset is shown in Figure S3. Cases, where models did not find all the top variants within 100 rounds, were not used to compute the averages.

### 9.6 Ability to Rank Mutants Correctly

The ability to rank samples correctly seems to be an important factor for selecting the next points for experimental validation. The model with the highest average Kendall correlation coefficient to rank samples correctly RF_ESM-2_EI required the least number of rounds to identify the top N sequences. This is shown in Figure S4.

The exact values measured for all models, representations and acquisition functions are listed in Table S3.

### 9.7 Model performance on different datasets

Model performances differ between datasets. The average number of rounds needed to discover the top variants for each model calls are shown in the Plots S5, S6, and S7.

Analysis of the plots reveals that some for some datasets, such as the CP2C9_HUMAN_Amorosi _2021_activity dataset or the OXDA_RHOTO _Vanella_2023_activity dataset it was consistently more difficult to discover the top proteins in the datasets for all the models, as indicated from the increased number of rounds needed to discover the top proteins.

### 9.8 Feature Importance for Identifying the Best Proteins

To understand the factors influencing a model’s ability to identify the top-performing proteins, we analyzed the importance of three features: Average Ranking Ability, Average Mean Squared Error (MSE), and Average Correlation. The goal was to determine how these features predict the model’s efficiency in discovering the top 5, 10, 20, and 50 proteins, where a lower number of rounds indicates better performance.

#### Methods

We employed two analytical approaches: correlation analysis and feature importance using RR.

1. **Correlation Analysis:** We computed Pearson correlation coefficients between the three features and the number of rounds required to identify the top proteins. This provided an initial understanding of the linear relationships between the features and model performance.
2. **RR Coefficients:** We utilized RR to predict the number of rounds required to identify the top 5, 10, 20, and 50 proteins, using the three features as predictors. RR provides coefficients for each feature, indicating the direction and magnitude of their relationship with the target variables.

#### Results

The correlation analysis revealed varying relationships between the features and the number of rounds required to identify the top proteins (Table S4). The RR model coefficients provided further insights into the linear relationship between each feature and the targets (Table S5).

Table S4 shows that Average Ranking Ability has a moderate negative correlation with the number of rounds required to identify the top 5 and 50 proteins, indicating that higher ranking ability may lead to faster identification of the best proteins. Average MSE exhibits a moderate positive correlation with the targets, suggesting that models with lower MSE tend to find the top proteins more quickly. Average Correlation shows weak correlations with the targets, indicating a less direct impact on the model’s performance.

**Figure S3:**
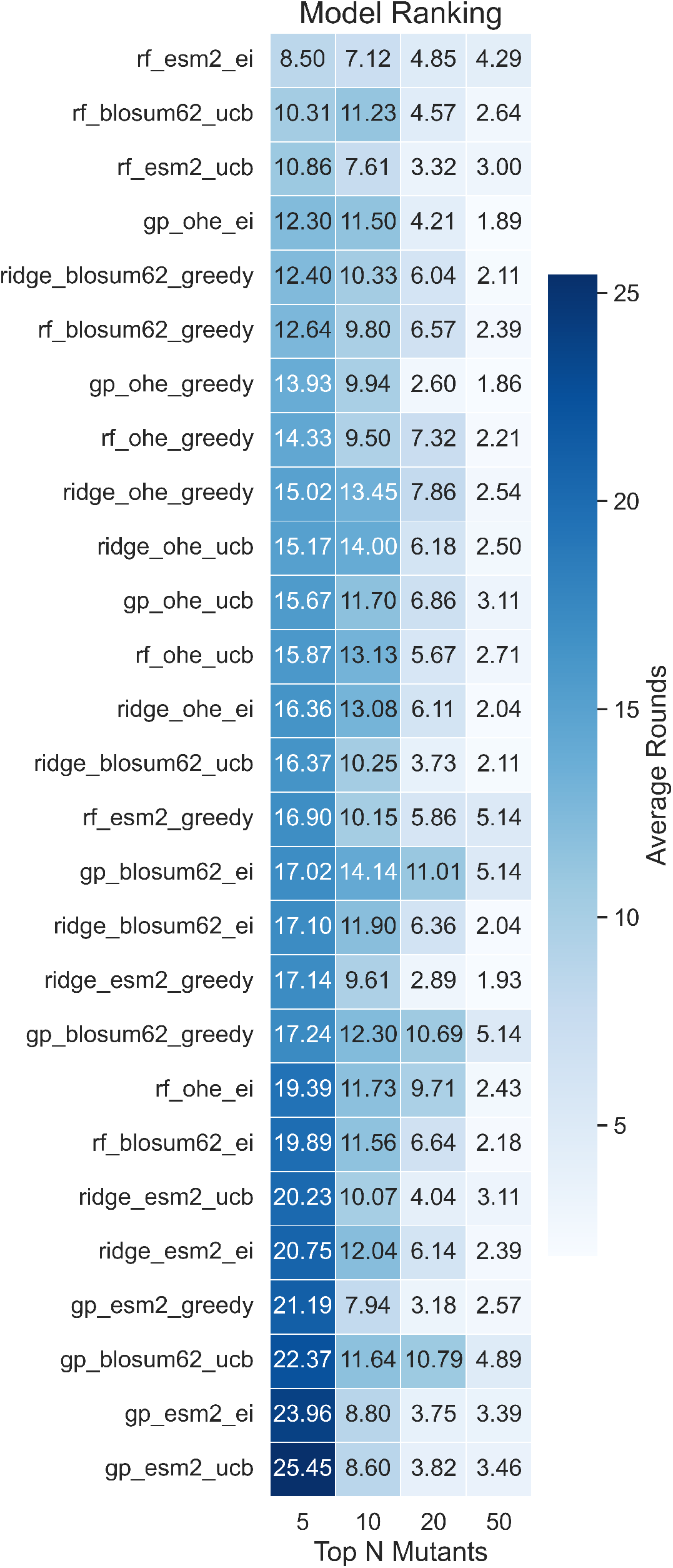
Average number of rounds needed to discover the top mutants in a dataset for each model.

**Figure S4:**
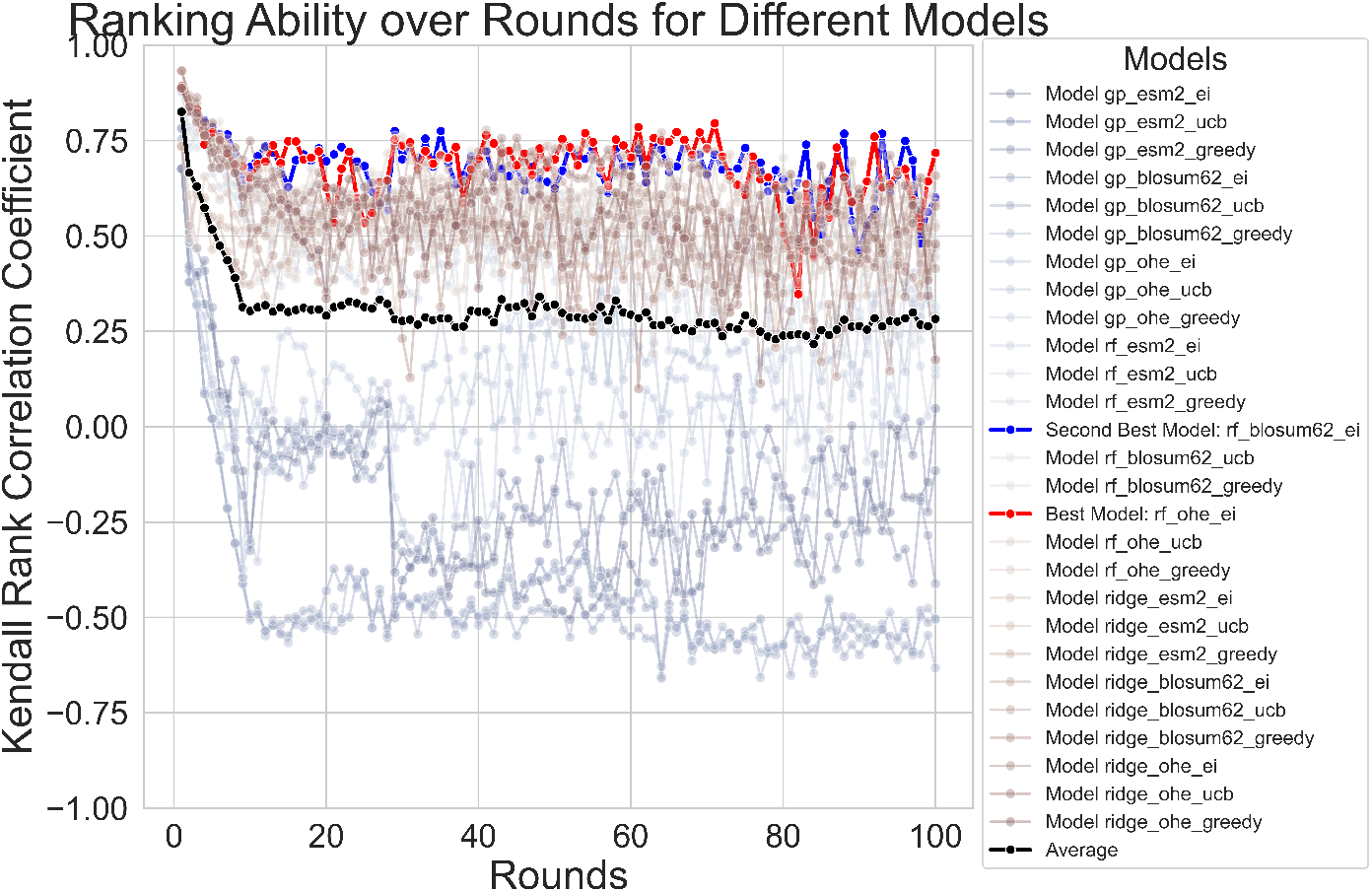
Ability of models to rank mutants correctly.

**Table S3:** Average correlation of different models.

| Models | Kendall Rank Correlation |
| --- | --- |
| RF_ESM-2_EI | 0.780855 |
| Ridge_ESM-2_EI | 0.682431 |
| Ridge_OHE_UCB | 0.648159 |
| RF_ESM-2_Greedy | 0.635948 |
| RF_OHE_EI | 0.631212 |
| Ridge_OHE_EI | 0.622729 |
| RF_BLOSUM62_EI | 0.605914 |
| Ridge_OHE | 0.605047 |
| Ridge_ESM-2_Greedy | 0.586710 |
| Ridge_BLOSUM62_EI | 0.579155 |
| Ridge_ESM-2_UCB | 0.571636 |
| RF_ESM-2_UCB | 0.540266 |
| GP_BLOSUM62_EI | 0.526180 |
| GP_BLOSUM62_Greedy | 0.524859 |
| GP_BLOSUM62_UCB | 0.518750 |
| Ridge_BLOSUM62_Greedy | 0.513484 |
| RF_OHE_Greedy | 0.502725 |
| Ridge_BLOSUM62_UCB | 0.500854 |
| GP_ESM-2_EI | 0.481600 |
| RF_BLOSUM62_UCB | 0.474556 |
| RF_BLOSUM62_Greedy | 0.472169 |
| GP_ESM-2_UCB | 0.446015 |
| GP_ESM-2_Greedy | 0.445125 |
| RF_OHE_UCB | 0.397548 |
| GP_OHE_Greedy | 0.328683 |
| GP_OHE_UCB | 0.269706 |
| GP_OHE_EI | 0.246086 |

**Table S4:** Pearson Correlation Coefficients between Features and Number of Rounds to Identify Top Proteins.

| Feature | Top 5 | Top 10 | Top 20 | Top 50 |
| --- | --- | --- | --- | --- |
| Average Ranking Ability | -0.395 | 0.041 | -0.261 | -0.603 |
| Average MSE | 0.196 | 0.176 | 0.343 | 0.494 |
| Average Correlation | 0.020 | 0.015 | 0.205 | 0.193 |

**Table S5:** RR coefficients for predicting number of rounds to identify top proteins.

| Feature | Top 5 | Top 10 | Top 20 | Top 50 |
| --- | --- | --- | --- | --- |
| Average Ranking Ability | -3.61777 | 1.30097 | -0.44659 | -1.17903 |
| Average MSE | -0.35112 | 1.00998 | 1.08761 | 0.30908 |
| Average Correlation | 0.76722 | 0.91358 | 2.00848 | 1.01540 |

**Figure S5:**
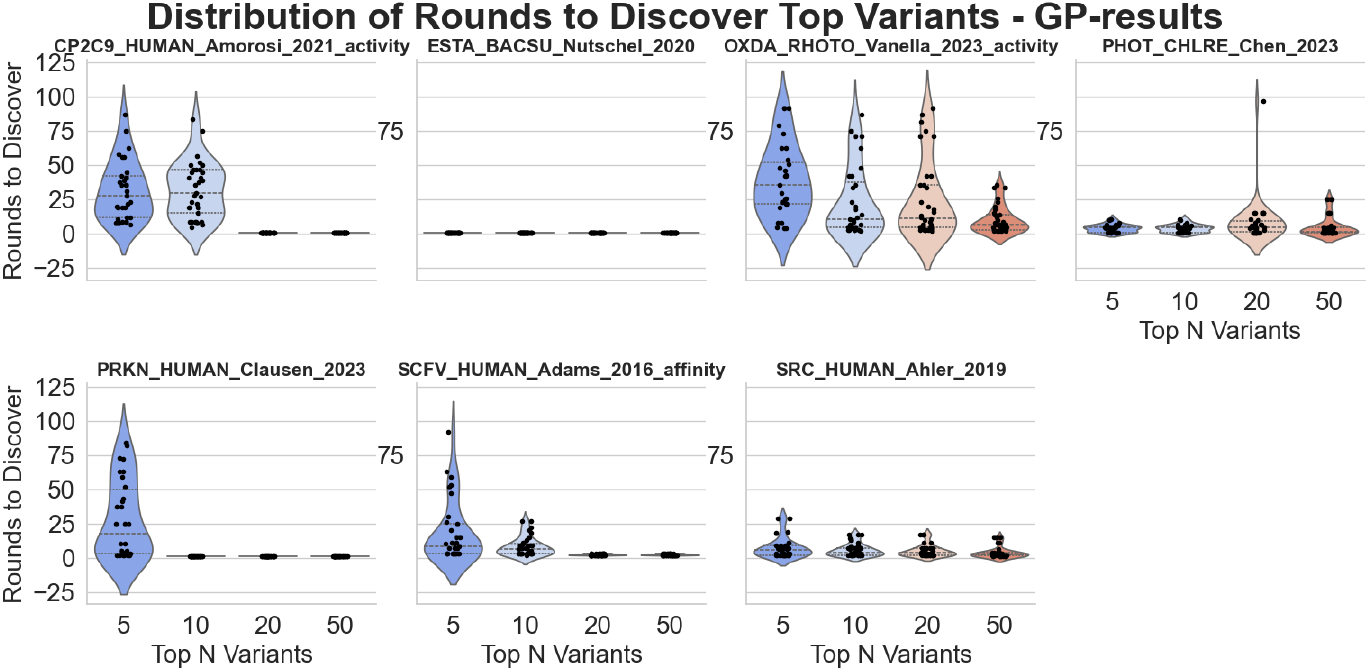
Performance of GP model on individual datasets.

**Figure S6:**
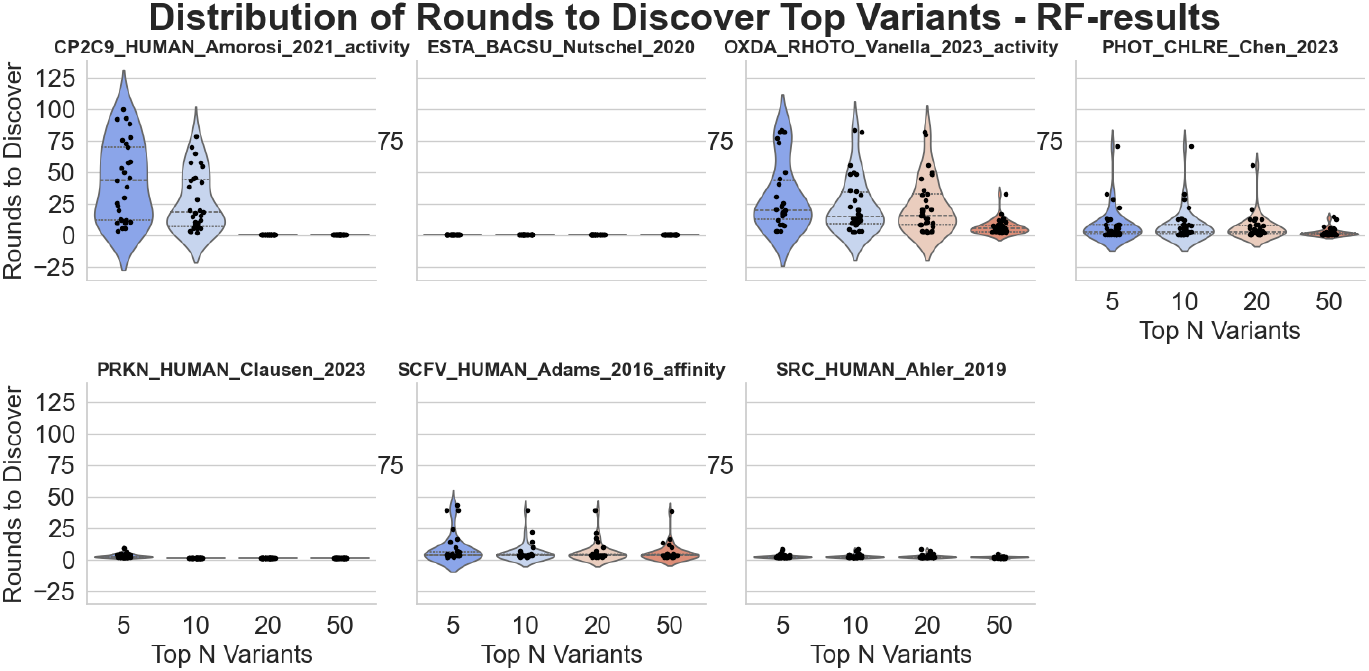
Performance of RF model on individual datasets.

**Figure S7:**
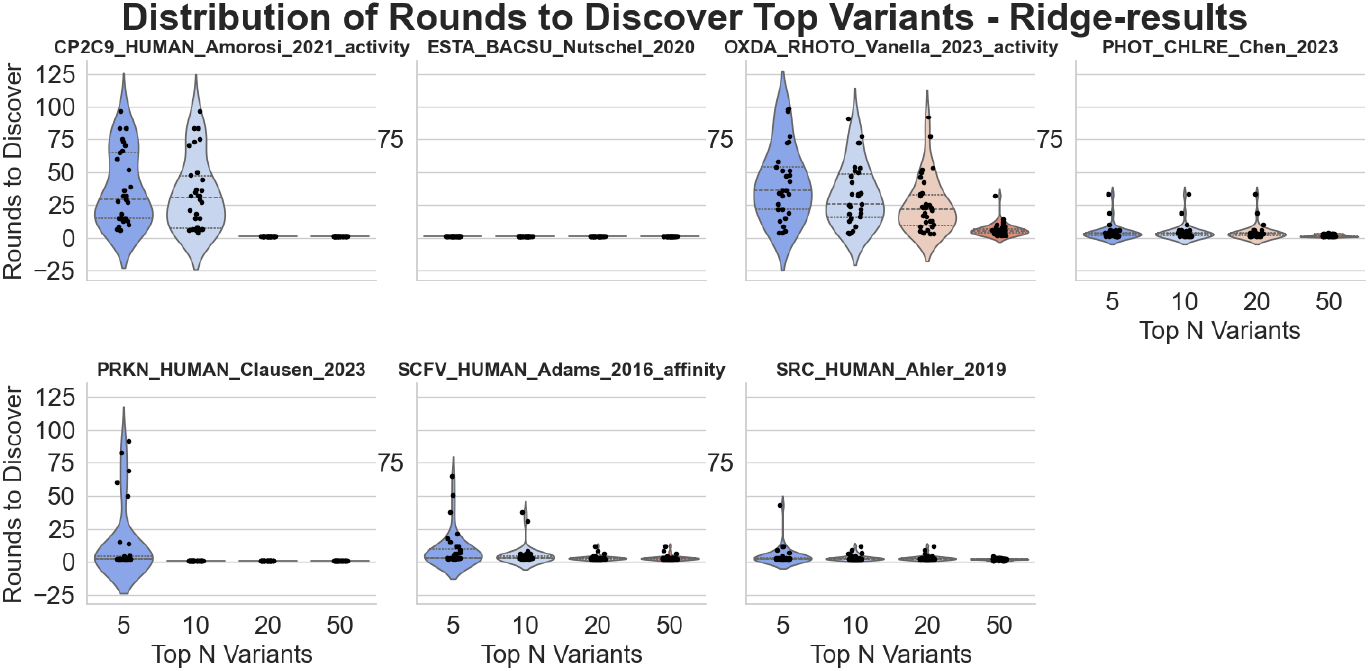
Performance of RR model on individual datasets.

Table S5 shows that ‘Average Ranking Ability’ has a strong negative impact on the number of rounds required to identify the top 5 proteins, suggesting that higher ranking ability leads to faster identification. Conversely, ‘Average MSE’ has a stronger positive effect on larger subsets, implying that models with lower MSEs tend to perform better across different target sizes. Lastly, ‘Average Correlation’ becomes increasingly important for identifying larger sets (e.g., top 20 and top 50), indicating that better uncertainty estimation is beneficial for finding more top-performing proteins. The recorded metrics for all models, representations, and acquisition functions are listed in Table S6.

### 9.9 Dataset Selection for Benchmark Studies

This study aimed to identify the top mutants in a dataset. Thus it was vital to ensure that the top variants in the uploaded dataset are not experimental artefacts or errors. Different datasets were selected to cover different properties. The reasons for their inclusion are summarized in Table S7.

The SCFV_HUMAN_Adams_2016_affinity is a binding dataset that is not sourced from the ProteinGym, but instead from the from an original publication by Adams et al. [59]. The dataset was created by extracting the sequences that were measured in all three replicate files provided on GitHub, and computing the average KD for them, used as fitness values. The sequences were reconstructed from the wild-type sequence and the available CDR1 and CDR3 regions.

**Table S6:**
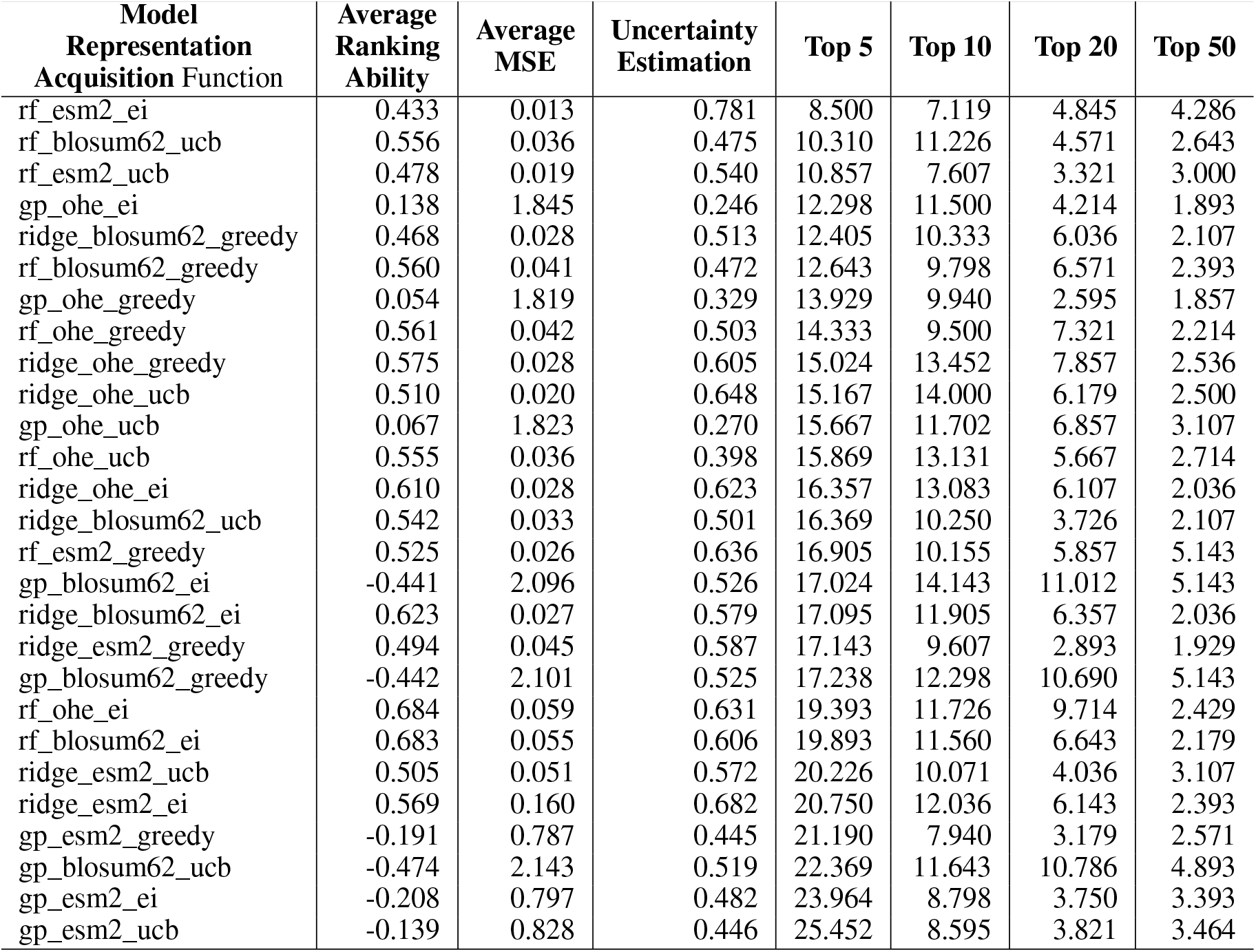
Tabular overview of model metrics, sorted by the number of rounds needed to identify top 5 proteins.

| Model Representation Acquisition Function | Average Ranking Ability | Average MSE | Uncertainty Estimation | Top 5 | Top 10 | Top 20 | Top 50 |
| --- | --- | --- | --- | --- | --- | --- | --- |
| rf_esm2_ei | 0.433 | 0.013 | 0.781 | 8.500 | 7.119 | 4.845 | 4.286 |
| rf_blosum62_ucb | 0.556 | 0.036 | 0.475 | 10.310 | 11.226 | 4.571 | 2.643 |
| rf_esm2_ucb | 0.478 | 0.019 | 0.540 | 10.857 | 7.607 | 3.321 | 3.000 |
| gp_ohe_ei | 0.138 | 1.845 | 0.246 | 12.298 | 11.500 | 4.214 | 1.893 |
| ridge_blosum62_greedy | 0.468 | 0.028 | 0.513 | 12.405 | 10.333 | 6.036 | 2.107 |
| rf_blosum62_greedy | 0.560 | 0.041 | 0.472 | 12.643 | 9.798 | 6.571 | 2.393 |
| gp_ohe_greedy | 0.054 | 1.819 | 0.329 | 13.929 | 9.940 | 2.595 | 1.857 |
| rf_ohe_greedy | 0.561 | 0.042 | 0.503 | 14.333 | 9.500 | 7.321 | 2.214 |
| ridge_ohe_greedy | 0.575 | 0.028 | 0.605 | 15.024 | 13.452 | 7.857 | 2.536 |
| ridge_ohe_ucb | 0.510 | 0.020 | 0.648 | 15.167 | 14.000 | 6.179 | 2.500 |
| gp_ohe_ucb | 0.067 | 1.823 | 0.270 | 15.667 | 11.702 | 6.857 | 3.107 |
| rf_ohe_ucb | 0.555 | 0.036 | 0.398 | 15.869 | 13.131 | 5.667 | 2.714 |
| ridge_ohe_ei | 0.610 | 0.028 | 0.623 | 16.357 | 13.083 | 6.107 | 2.036 |
| ridge_blosum62_ucb | 0.542 | 0.033 | 0.501 | 16.369 | 10.250 | 3.726 | 2.107 |
| rf_esm2_greedy | 0.525 | 0.026 | 0.636 | 16.905 | 10.155 | 5.857 | 5.143 |
| gp_blosum62_ei | -0.441 | 2.096 | 0.526 | 17.024 | 14.143 | 11.012 | 5.143 |
| ridge_blosum62_ei | 0.623 | 0.027 | 0.579 | 17.095 | 11.905 | 6.357 | 2.036 |
| ridge_esm2_greedy | 0.494 | 0.045 | 0.587 | 17.143 | 9.607 | 2.893 | 1.929 |
| gp_blosum62_greedy | -0.442 | 2.101 | 0.525 | 17.238 | 12.298 | 10.690 | 5.143 |
| rf_ohe_ei | 0.684 | 0.059 | 0.631 | 19.393 | 11.726 | 9.714 | 2.429 |
| rf_blosum62_ei | 0.683 | 0.055 | 0.606 | 19.893 | 11.560 | 6.643 | 2.179 |
| ridge_esm2_ucb | 0.505 | 0.051 | 0.572 | 20.226 | 10.071 | 4.036 | 3.107 |
| ridge_esm2_ei | 0.569 | 0.160 | 0.682 | 20.750 | 12.036 | 6.143 | 2.393 |
| gp_esm2_greedy | -0.191 | 0.787 | 0.445 | 21.190 | 7.940 | 3.179 | 2.571 |
| gp_blosum62_ucb | -0.474 | 2.143 | 0.519 | 22.369 | 11.643 | 10.786 | 4.893 |
| gp_esm2_ei | -0.208 | 0.797 | 0.482 | 23.964 | 8.798 | 3.750 | 3.393 |
| gp_esm2_ucb | -0.139 | 0.828 | 0.446 | 25.452 | 8.595 | 3.821 | 3.464 |

**Table S7:** ProteinGym datasets selected for the benchmark with summarized reasons for inclusion.

| Dataset mutant? | Multi Property | Comments |  |
| --- | --- | --- | --- |
| CP2C9_HUMAN_Amorosi_2021_activity | no | activity | Assay quality looks good (figures 1e and 2). Score distribution is unusual, but log-transforming the data may correct behavior during sampling. |
| ESTA_BACSU_Nutschel_2020 | no | thermostability | High-quality dataset, likely better than DMS studies. WT temperature is around 45°C. |
| OXDA_RHOTO_Vanella_2023_activity | yes | activity | Good quality (fig 2G). Data shows good $R^2$. |
| PHOT_CHLRE_Chen_2023 | no | fluorescence | High $R^2$ value between pooled and individual measurements (fig 1C). |
| PRKN_HUMAN_Clausen_2023 | no | stability | Benchmark shows dataset performs well (referenced to fig 4f). |
| SRC_HUMAN_Ahler_2019 | no | activity | High-quality assay (fig 2I). |
| SCFV_HUMAN_Adams_2016_affinity | yes | binding affinity | High-quality assay (fig. 4C). Has not been sourced from the ProteinGym. |

**Table S8:**
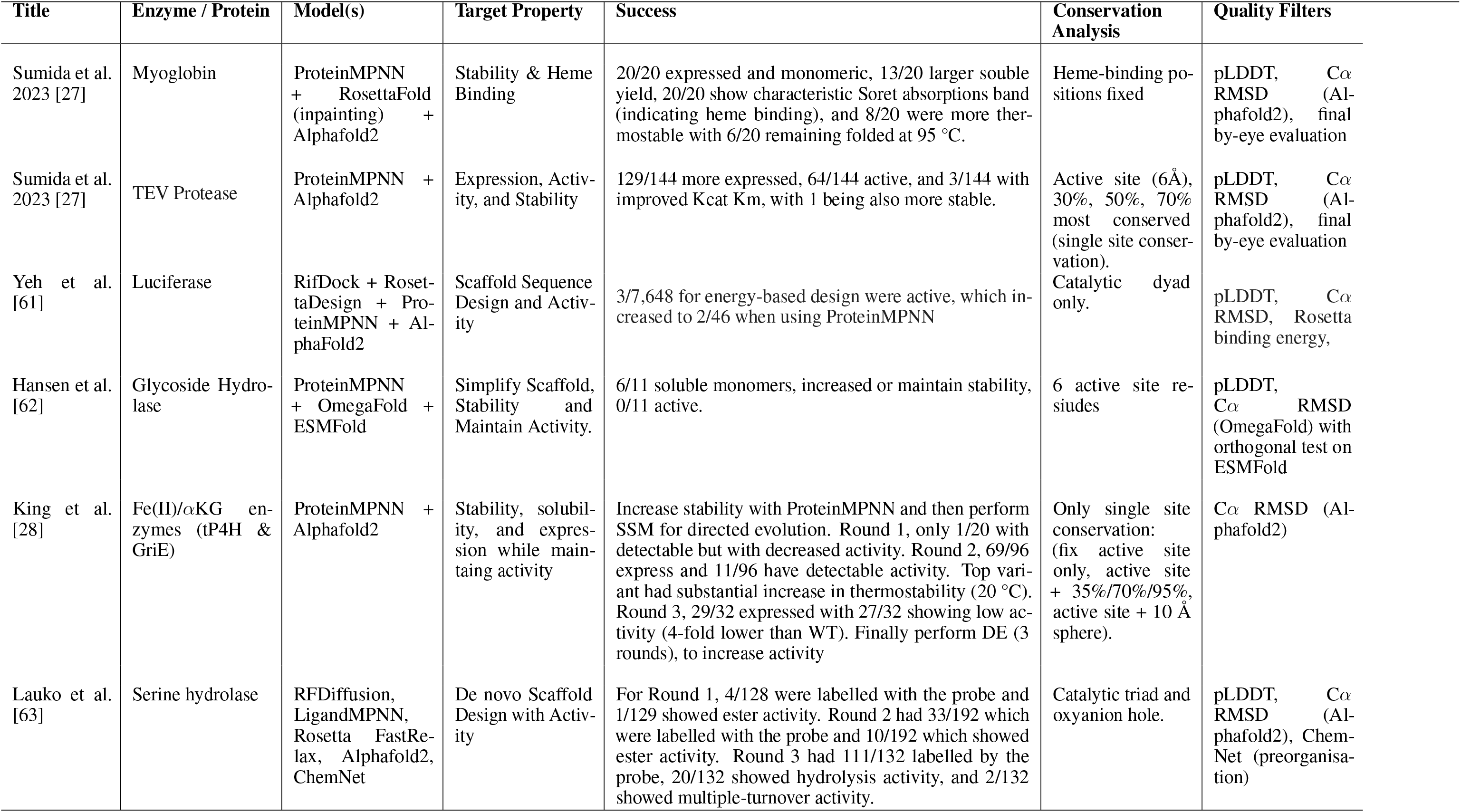
Overview of Protein Design Applications using Inverse Folding Algorithms. For several protein design protocols, combining several orthogonal methods, the table provides and overview of success, quality filters, and otherwise design considerations which may have increased the hit-rate.

**Table S9:**
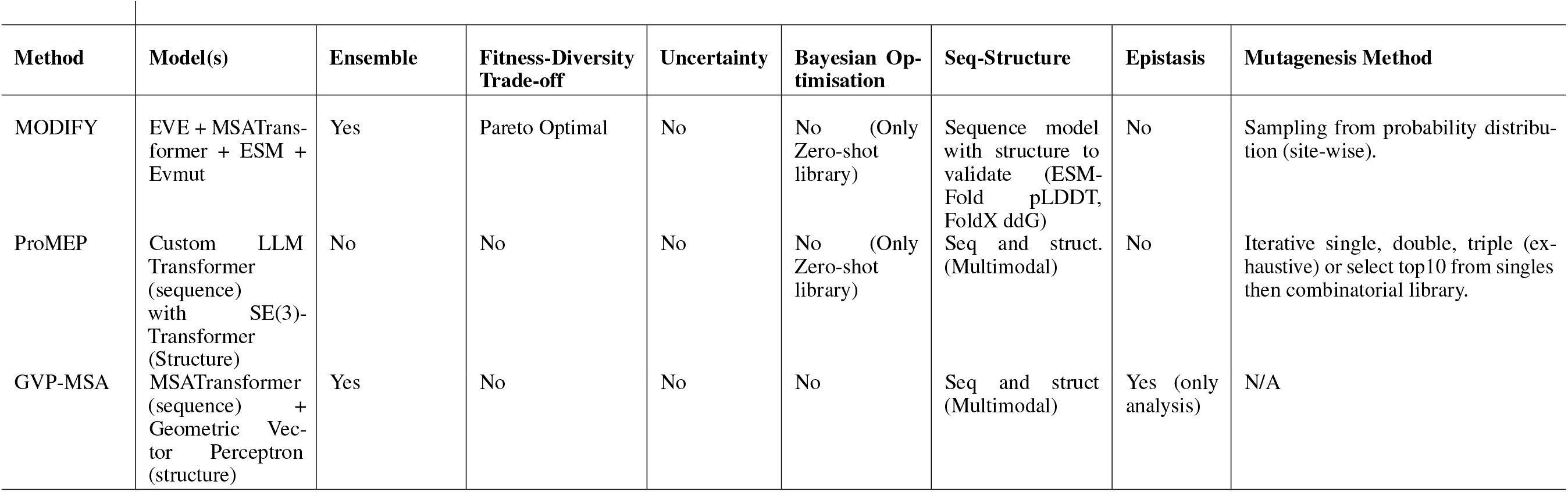
Summary of SOTA methods for Protein Engineering and Design. Only MODIFY addresses Pareto optimial fronteirs to balance sequence diversity and mean-fitness of the generated library. All three methods, notably, do not use uncertainty in the fitness prediction and likewise are not built for Bayesian optimisation. MODIFY and ProMEP do build zero-shot libraries. None of the models directly model epistatic effects. Interestingly, none of the methods implement genetic algorithms to build new sequences for the next library. MODIFY samples only based on site-wise probabilities for each amino acid, thus neglecting co-evolving pairs, while ProMEP brute forces exploration by procuring fitness predictions for all single/double variants.

